# Which animals are at risk? Predicting species susceptibility to Covid-19

**DOI:** 10.1101/2020.07.09.194563

**Authors:** MR Alexander, CT Schoeder, JA Brown, CD Smart, C Moth, JP Wikswo, JA Capra, J Meiler, W Chen, MS Madhur

## Abstract

In only a few months, the novel coronavirus severe acute respiratory syndrome coronavirus 2 (SARS-CoV-2) has caused a global pandemic, leaving physicians, scientists, and public health officials racing to understand, treat, and contain this zoonotic disease. SARS-CoV-2 has made the leap from animals to humans, but little is known about variations in species susceptibility that could identify potential reservoir species, animal models, and the risk to pets, wildlife, and livestock. While there is evidence that certain species, such as cats, are susceptible, the vast majority of animal species, including those in close contact with humans, have unknown susceptibility. Hence, methods to predict their infection risk are urgently needed. SARS-CoV-2 spike protein binding to angiotensin converting enzyme 2 (ACE2) is critical for viral cell entry and infection. Here we identified key ACE2 residues that distinguish susceptible from resistant species using in-depth sequence and structural analyses of ACE2 and its binding to SARS-CoV-2. Our findings have important implications for identification of ACE2 and SARS-CoV-2 residues for therapeutic targeting and identification of animal species with increased susceptibility for infection on which to focus research and protection measures for environmental and public health.

## INTRODUCTION

Severe acute respiratory syndrome coronavirus 2 (SARS-CoV-2) is the virus responsible for the global pandemic of coronavirus disease-2019 (Covid-19) that is impacting millions of lives and the global economy. Covid-19 is a zoonotic infection capable of crossing the species barrier. SARS-CoV-2 is thought to have originated in bats and subsequently transmitted to humans, perhaps through a secondary host.^1,2^ Emerging experimental and observational evidence demonstrates differences in species susceptibility to infection. For example, humans, house cats, tigers, and lions are all susceptible to infection by SARS-CoV-2.^3-6^ Golden Syrian hamsters and rhesus monkeys are also capable of being experimentally infected by SARS-CoV-2 and developing Covid-19 pathologies.^7,8^ In contrast, observational and experimental studies with direct intranasal inoculation have demonstrated that chickens, ducks, and mice are not susceptible to SARS-CoV-2 infection.^5,9-11^ Interestingly however, susceptibility is not dichotomous. Although ferrets are also susceptible to infection, intranasal inoculation failed to result in spread of infection to the lower respiratory tract, significantly limiting symptom development.^5^ In addition, although dogs failed to exhibit infection of the respiratory tract and appear asymptomatic, a minority of experimentally or environmentally exposed dogs exhibited evidence of infection by SARS-CoV-2 PCR or SARS-CoV-2 seroconversion with production of SARS-CoV-2-specific antibodies.^5,12^ While pigs have not demonstrated evidence of infection after intranasal inoculation, overexpression of swine ACE2 in cultured cells supports some degree of viral entry.^5,9,13^ Hence, ferrets, dogs, and pigs are classified as having intermediate susceptibility to infection. Despite these findings, the number of animal species tested for susceptibility to infection in experimental or observational studies is very limited. Thus, methods of determining risk of species with unknown susceptibility are urgently needed to reduce risk of propagating transmission, protect food supplies, identify potential intermediate hosts, and discover animal models for research. Identifying the key residues mediating susceptibility to infection can also guide rational drug design.

SARS-CoV-2 is a member of the coronavirus family of single-stranded RNA viruses.^9^ The spike protein on the surface of the SARS-CoV-2 virus mediates interaction with its receptor, angiotensin converting enzyme 2 (ACE2), to promote membrane fusion and virus entry into the cell. The receptor binding domain (RBD) of the spike protein contains a receptor binding motif (RBM) that binds to the peptidase domain of ACE2.^14^ Following spike protein cleavage, fusion of the viral and host cell membranes occurs to enable viral entry into the cell.^15^ Interaction of the SARS-CoV-2 spike protein RBD and ACE2 is thus critical for viral cell entry and infection.^9^ The importance of this interaction in infection is further supported by evidence that exogenous soluble ACE2 limits infection in human organoids,^10^ and that overexpression of human ACE2 is necessary to enable viral cell entry in HeLa cells *in vitro* and SARS-CoV-2 infection in mouse models *in vivo*.^9,16^

ACE2 is present in almost all vertebrates, however sequence differences exist that may hold clues to differences in SARS-CoV-2 susceptibility, as has been observed for SARS-CoV.^17,18^ Understanding such differences could provide insight into key structural interactions between ACE2 and SARS-CoV-2 RBD important for infection, and permit development of a susceptibility score for estimating the infection risk of various species. In this manuscript we integrate experimentally validated differences in susceptibility to SARS-CoV-2 infection with ACE2 sequence comparisons and in-depth structural analyses to determine how differences in ACE2 across species influence interaction with SARS-CoV-2 RBD. We identified multiple key residues mediating structural interactions between ACE2 and SARS-CoV-2 RBD and use these residues to generate a susceptibility score to predict animals with elevated risk of infection. We also demonstrate that SARS-CoV-2 is nearly optimal for binding ACE2 of humans compared to other animals, which may underlie the highly contagious nature of this virus amongst humans. Our findings have important implications for identification of ACE2 and SARS-CoV-2 residues for therapeutic targeting and identification of animal species with increased susceptibility for infection on which to focus research and protection efforts.

## RESULTS

### Susceptibility does not segregate according to phylogeny and ACE2 sequence similarity

Given experimental evidence for susceptibility of humans, house cats, tigers, lions, rhesus macaques, and Golden Syrian hamsters to SARS-CoV-2 infection, and experimental evidence for non-susceptibility of mice, ducks, and chickens,^3-5,7,9-11,19^ we performed protein sequence alignment of ACE2 from these organisms using MAFFT (**Extended Data Figure 1**).^20^ We also included species with intermediate susceptibility, including dogs, pigs, and ferrets,^5,9,12^ as well as species with unknown susceptibility, including camels, horses, Malayan pangolin, and sheep. The degree of similarity of ACE2 protein sequences largely fell along expected phylogenetic relationships among species (**Extended Data Figure 2**). Susceptibility to SARS-CoV-2 infection, however, did not match either phylogenetic relationships or ACE2 sequence similarities across species. For example, mouse (*Mus musculus*) is not susceptible to infection. However, mouse ACE2 sequence is more similar to a susceptible species, Golden Syrian hamster (*Mesocricetus auratus*), than non-susceptible species such as duck (*Aythya fuligula*) or chicken (*Gallus gallus*).^9,16^ In addition, mice are phylogenetically more similar to susceptible species such as humans (*Homo sapiens*) and rhesus macaques (*Macaca mulatta*) than non-susceptible species such as ducks and chicken.^9,16^ These findings suggest that neither phylogenetic relationships nor overall ACE2 protein sequence similarity across species is able to predict susceptibility to SARS-CoV-2 infection.

### Sequence alignment identifies ACE2 residues distinguishing susceptible from non-susceptible species

An alternative approach is to use the experimentally validated differences in infection susceptibility across species to focus on ACE2 amino acids that most differ between susceptible and non-susceptible species. We thus calculated a weighted score of how well the aligned amino acids stratify susceptible versus non-susceptible species, incorporating amino acid similarity. This score, termed GroupSim, permits quantitative determination of which amino acids in the alignment best stratify susceptible from non-susceptible species.^21^ This analysis demonstrated that multiple amino acid positions in the ACE2 alignment, including Leu79, His34, Tyr83, and Gln24, are highly similar in susceptible species and quite different in non-susceptible species (**Extended Data Table 1 and Supplemental Table 1**). When mapping these scores onto the structure of the SARS-CoV-2 RBD and ACE2 complex, multiple residues with high GroupSim scores were present at or near the binding interface including His34, Asp30, Thr92, Gln24, Lys31, and Leu79 (**Figure 1**). We then extended this analysis by focusing on key residues previously demonstrated from prior structural analysis to be important for ACE2 and SARS-CoV-2 RBD interactions (**Table 1**).^7,22-24^ Interestingly, this revealed that key amino acids for the ACE2 and SARS-CoV-2 spike protein interaction were enriched among the top scoring GroupSim positions (7 of 35; p<0.0001; Fisher’s exact test). Such key residues based on structural analysis being over-represented in amino acid positions that best discriminated susceptible from non-susceptible species suggests that structural interactions between ACE2 and SARS-CoV-2 spike protein importantly determine differences in species susceptibility to infection. In addition, these data suggest that certain ACE2 amino acid residues may be particularly important for determining susceptibility, including Leu79, His34, Tyr83, Gln24, Lys31, Asp30, and Glu329.

**Table 1:** Twenty-four key residues for SARS-CoV-2 RBD and ACE2 interactions.

| Genus species | Common name | 19 | 24 | 27 | 28 | 30 | 31 | 34 | 35 | 37 | 38 | 41 | 42 | 45 | 79 | 82 | 83 | 325 | 329 | 330 | 353 | 354 | 355 | 357 | 393 |
| --- | --- | --- | --- | --- | --- | --- | --- | --- | --- | --- | --- | --- | --- | --- | --- | --- | --- | --- | --- | --- | --- | --- | --- | --- | --- |
| *Homo sapiens* | Human | S | Q | T | F | D | K | H | E | E | D | Y | Q | L | L | M | Y | Q | E | N | K | G | D | R | R |
| *Macaca mulatta* | Rhesus macaque | S | Q | T | F | D | K | H | E | E | D | Y | Q | L | L | M | Y | Q | E | N | K | G | D | R | R |
| *Felis catus* | House cat | S | L | T | F | E | K | H | E | E | E | Y | Q | L | L | T | Y | Q | E | N | K | G | D | R | R |
| *Panthera tigris altaica* | Tiger | S | L | T | F | E | K | H | E | E | E | Y | Q | L | L | T | Y | Q | E | N | K | G | D | R | R |
| *Panthera leo* | Lion | S | L | T | F | E | K | H | E | E | E | Y | Q | L | L | T | Y | Q | E | N | K | G | D | R | R |
| *Mesocricetus auratus* | Golden Syrian hamster | S | Q | T | F | D | K | Q | E | E | D | Y | Q | L | L | N | Y | Q | E | N | K | G | D | R | R |
| *Mus musculus* | Mouse | S | N | T | F | N | N | Q | E | E | D | Y | Q | L | T | S | F | Q | A | N | H | G | D | R | R |
| *Aythya fuligula* | Duck | D | - | M | F | A | E | V | R | E | D | Y | E | L | N | N | F | E | K | N | K | N | D | R | R |
| *Gallus gallus* | Chicken | D | - | T | F | A | E | V | R | E | D | Y | E | L | N | R | F | E | T | N | K | N | D | R | R |
| *Mustela putorius furo* | Ferret | S | L | T | F | E | K | Y | E | E | E | Y | Q | L | H | T | Y | E | Q | N | K | R | D | R | R |
| *Sus scrofa* | Pig | S | L | T | F | E | K | L | E | E | D | Y | Q | L | I | T | Y | Q | N | N | K | G | D | R | R |
| *Canis lupus familiaris* | Dog | S | L | T | F | E | K | Y | E | E | E | Y | Q | L | L | T | Y | Q | G | N | K | G | D | R | R |
| *Rhinolophus sinicus* | Chinese horseshoe bat | S | E | T | F | D | K | T | K | E | D | H | Q | L | L | N | Y | E | N | N | K | G | D | R | R |
| *Equus caballus* | Horse | S | L | T | F | E | K | S | E | E | E | H | Q | L | L | T | Y | Q | E | N | K | G | D | R | R |
| *Bos taurus* | Cow | S | Q | T | F | E | K | H | E | E | D | Y | Q | L | M | T | Y | Q | D | N | K | G | D | R | R |
| *Manis javanica* | Malayan pangolin | S | E | T | F | E | K | S | E | E | E | Y | Q | L | I | N | Y | Q | E | N | K | H | D | R | R |
| *Capra hircus* | Goat | S | Q | T | F | E | K | H | E | E | D | Y | Q | L | M | T | Y | Q | N | N | K | G | D | R | R |
| *Ovis aries* | Sheep | S | Q | T | F | E | K | H | E | E | D | Y | Q | L | M | T | Y | Q | D | N | K | G | D | R | R |
| *Camelus dromedarius* | Arabian Camel | S | L | T | F | E | E | H | E | E | D | Y | Q | L | T | T | Y | Q | D | N | K | G | D | R | R |
| *Camelus bactrianus* | Bactrian Camel | S | L | T | F | E | E | H | E | E | D | Y | Q | L | T | T | Y | Q | D | N | K | G | D | R | R |

**Figure 1:**
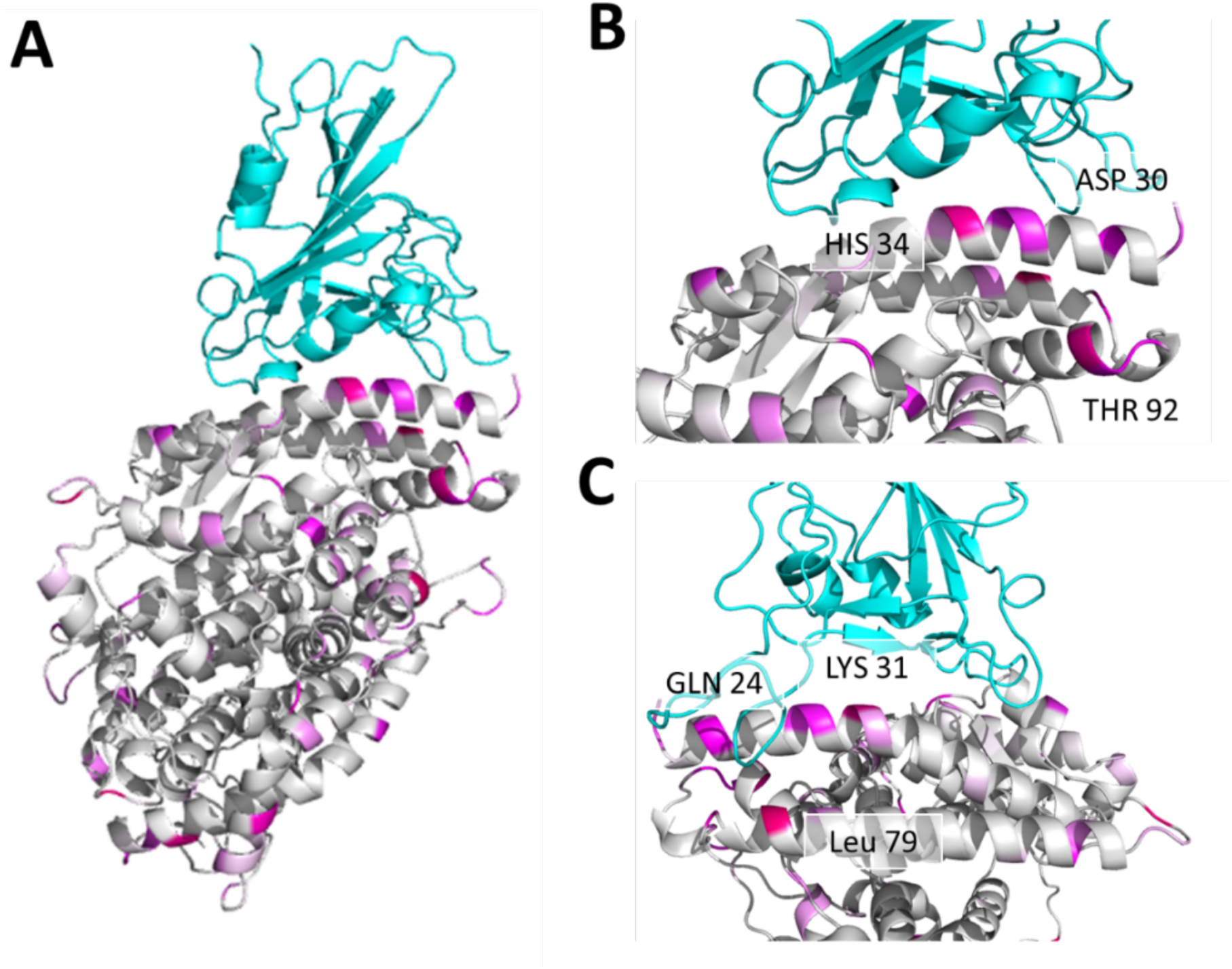
Multiple residues with high GroupSim scores are present at the interaction interface of the SARS-CoV-2 RBD and ACE2 complex. (**A**) SARS-CoV-2 RBD (top) and human ACE2 (bottom) complex shown as a ribbon diagram with GroupSim scores color coded in magenta. Higher scores are brighter in color. (**B**) Close-up view of the interface highlighting ACE2 residues with high GroupSim scores. (**C**) Close-up view after 90 degree rotation from (B) demonstrating additional residues at the interface with high GroupSim scores.

### SARS-CoV-2 has lower predicted binding affinity for ACE2 from non-susceptible avian species

We used homology modeling to identify structural determinants of binding the ACE2 protein from species with known differences in susceptibility to SARS-CoV-2 infection. The models were based on previously reported crystal structures of the human ACE2 in complex with SARS-CoV-2 (PDB: 6LZG and 6M0J).^14^ We modeled ACE2 in the presence of the SARS-CoV-2 RBD to allow backbone adjustment to the binder and refined by redocking of the RBD domain to optimize sidechains. Models were selected by overall calculated protein stability of the SARS-CoV-2 RBD complex, predicted binding energy between ACE2 and SARS-CoV-2 RBD, and similarity (as Cα-root mean square deviation [Cα-RMSD], **Extended Data Figure 3 and Extended Data Figure 4**). Based on these models, multiple approaches where undertaken to investigate the structural interactions between SARS-CoV-2-RBD and ACE2.

We evaluated the overall calculated protein stability and predicted binding energy for SARS-CoV-2-RBD and ACE2 complexes for each species. We considered the 100 best models for each species and evaluated evidence for difference in binding energy or stability between susceptible and non-susceptible species. The average mean predicted binding energy and calculated protein stability differs across species (**Figure 2**). Consistent with the lack of susceptibility of chickens (*Gallus gallus)*, chicken ACE2 in complex with SARS-CoV-2-RBD was the lowest scoring, or most energetically unfavorable model. The complex with duck ACE2 (*Aythya fuligula*) shows similarly unfavorable scores, indicating that ACE2 sequence differences leading to a lower structural binding ability in these two avian species may explain their lack of susceptibility to SARS-CoV-2 infection. However, the complex of SARS-CoV-2-RBD and ACE2 of the non-susceptible mouse (*Mus musculus*) exhibits lower binding energy and higher protein stability than several species that are susceptible, including the lion (*Panthera leo*), tiger (*Panthera tigris*), and cat (*Felis catus*). Thus, differences in SARS-CoV-2 and ACE2 complex stability have some discriminative power but are not the sole factor in differences in susceptibility across species.

**Figure 2:**
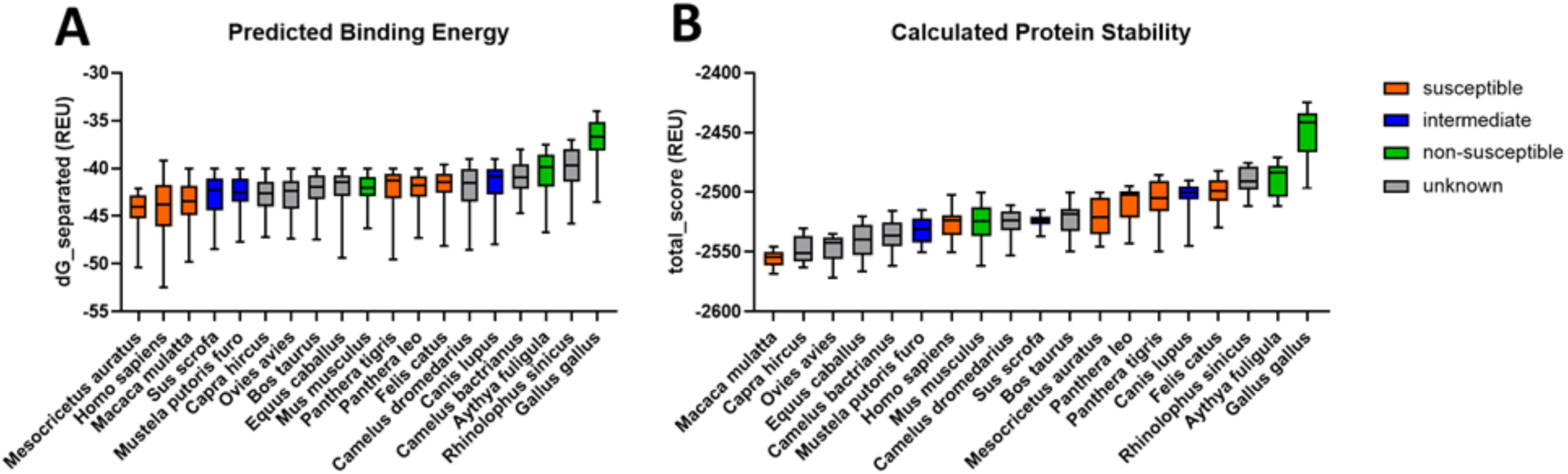
SARS-CoV-2 RBD has lower predicted binding energy and protein complex stability for ACE2 from non-susceptible avian species. (**A**) Predicted binding energy as calculated with Rosetta and (**B**) protein complex stability of SARS-CoV-2 RBD and ACE2 of various species predicted by Rosetta.

### Homology modeling identifies a link between ACE2 D30 and Y83 and SARS-CoV-2 susceptibility

As a complementary approach to determine whether particular residues may discriminate susceptible from non-susceptible species, we performed energetic modeling of residue-residue interactions in the interface of SARS-CoV-2 and ACE2 using Rosetta. Although the overall interaction pattern across residues is similar between susceptible, non-susceptible, and intermediate susceptibility species, there are significant differences in the magnitude of residue-residue interactions (**Figure 3**). For example, residue 30 (which is an aspartate in all susceptible species) forms a strong ionic interaction with lysine 417 of SARS-CoV-2 RBD and interacts modestly with other residues, including Phe456 and Tyr473. In contrast, in non-susceptible species such as chicken and duck where residue 30 contains an alanine this interaction is no longer present and is not substituted by any other structural rearrangements that might accommodate this change. Mouse (*Mus musculus)* ACE2 contains an asparagine in position 30 instead of an aspartate, which results in lower predicted binding energy due to the lack of an ionic interaction. A close-up view of residue 30 shows the different structural environment available in the non-susceptible species chicken, duck, and mouse as compared to susceptible species, including human (**Figure 4**). This analysis also identifies residue 83 of ACE2 as having differential energetic interactions across species. Residue 83 is a tyrosine in susceptible species and a phenylalanine in non-susceptible species (**Table 1**). Compared to susceptible species, this position exhibits significantly decreased binding energy with residues Asn487 and Tyr489 in SARS-CoV-2 RBD in non-susceptible species (**Figure 3**). Although ACE2 residue 83 also interacts with SARS-CoV-2 RBD phenylalanine 486, this interaction is unlikely to be significantly affected by differences between tyrosine and phenylalanine. However, the hydroxyl group of tyrosine at position 83 forms a hydrogen bond with the backbone oxygen of asparagine 487 that is negatively impacted by substitution to phenylalanine in non-susceptible species (**Figure 5A**). In addition to this residue-residue structural analysis, both ACE2 positions 30 and 83 were identified through the GroupSim analysis described above to be top residues discriminating susceptible from non-susceptible species based on sequence alignment (**Extended Data Table 1**). These results suggest that these amino acid positions of ACE2 may be important mediators of the structural interaction of ACE2 and SARS-CoV-2 RBD and determinants of differences to susceptibility to infection across species.

**Figure 3:**
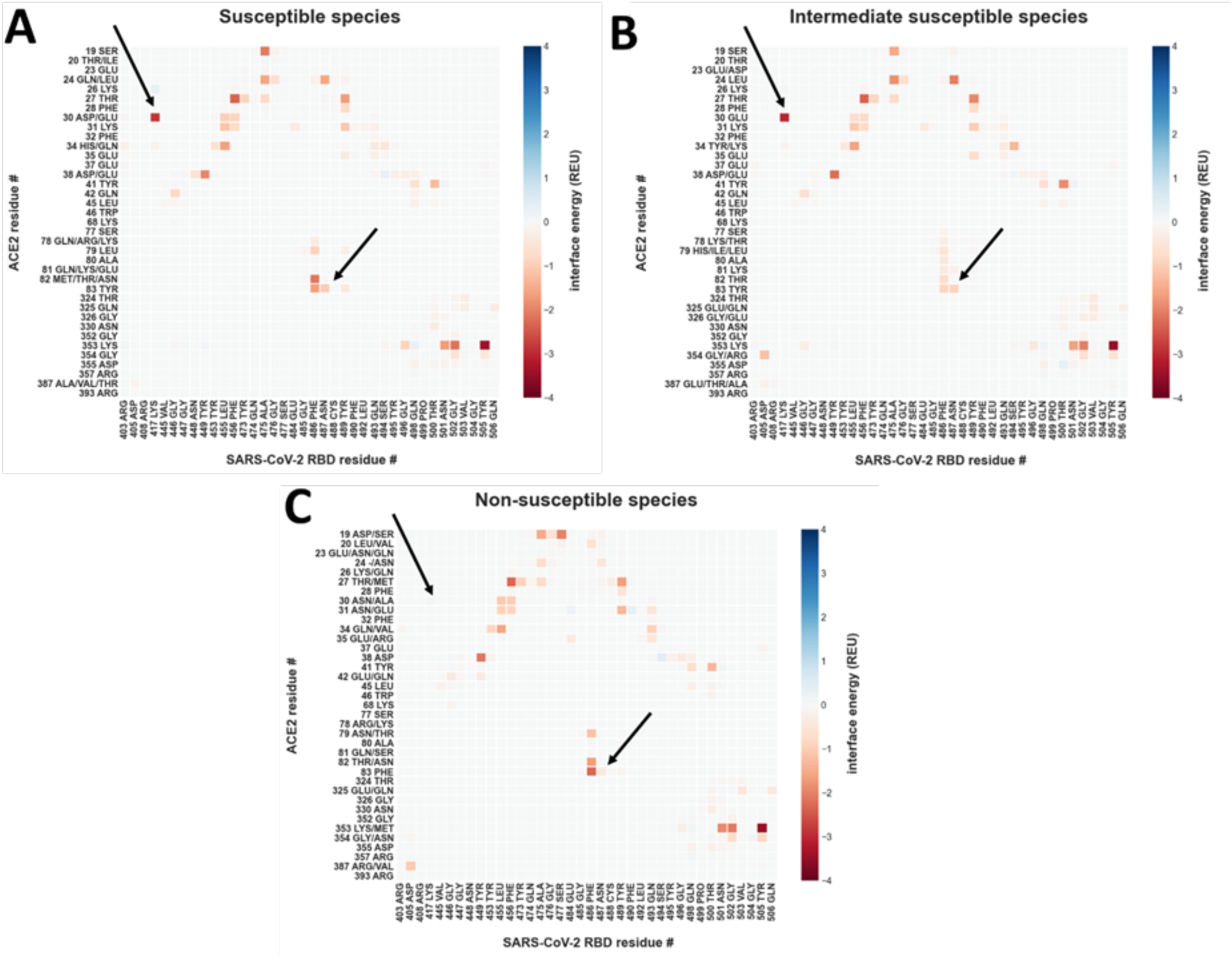
Energetic modeling of residue-residue interactions identifies a link between ACE2 D30 and Y83 and SARS-CoV-2 susceptibility. Residue-residue interactions are calculated with Rosetta, using the co-crystal structure of the human ACE2 in complex with the SARS-CoV-2-RBD (PDB: 6LZG and 6M0J) after backbone-constrained relaxation for all interactions greater than 0.05 Rosetta Energy Units (REU) or smaller than -0.05 REU. Interactions are presented as mean for all included samples. Residues depicted on the y-axis are all observed amino acid identities for the particular position in its susceptibility group. (**A**) Per-residue interactions for (A) susceptible species (human, cat, lion, tiger, hamster and rhesus macaque), (**B**) intermediate susceptibility species (pig, dog and ferret), and (**C**) non-susceptible species (duck, mouse, and chicken). The arrows point to interactions that are not observed in non-susceptible species.

**Figure 4:**
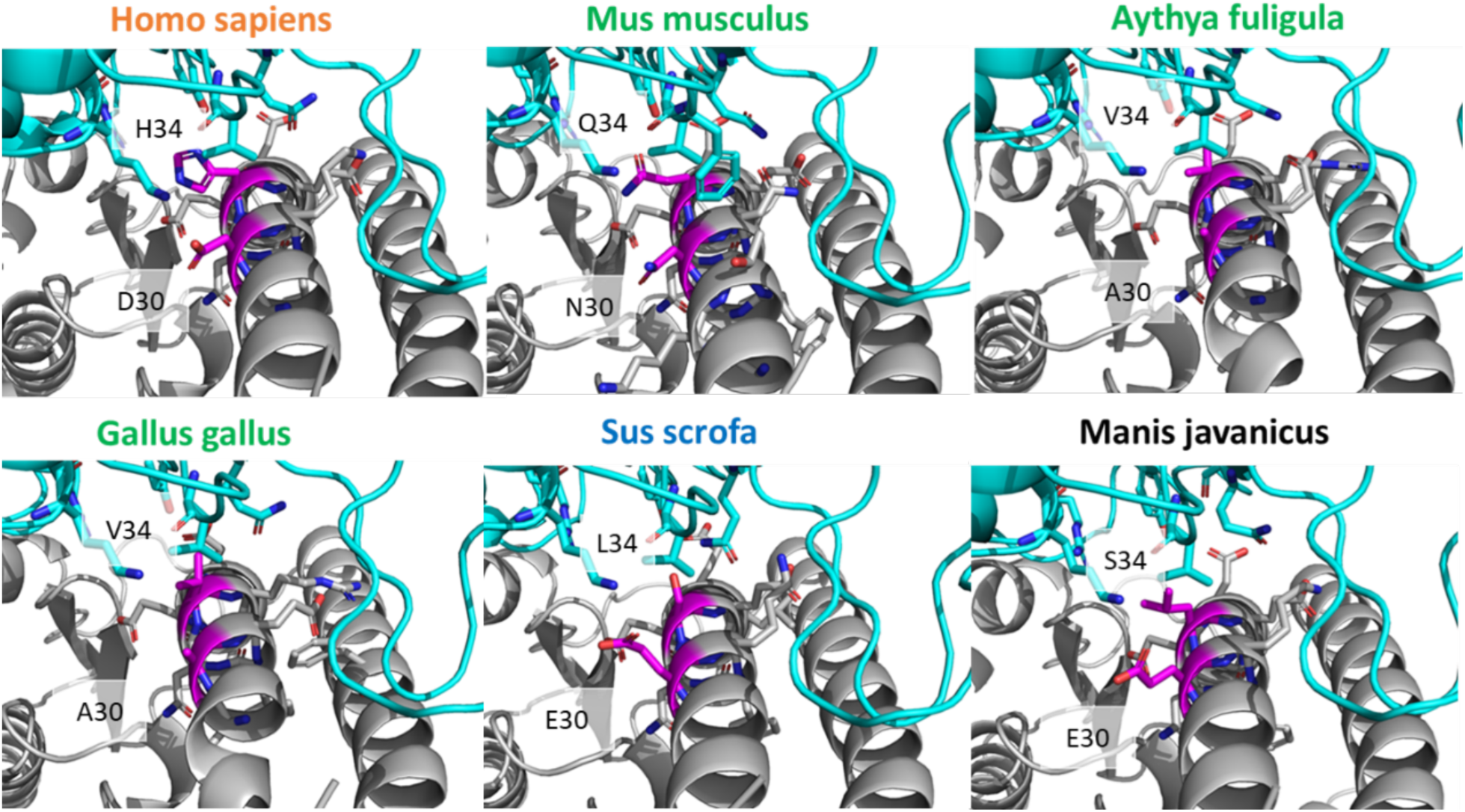
Binding interactions of ACE2 position 30 differ across species. Close-up of the differences in binding interactions of positions 30 and 34 (magenta) of ACE2 from each species with the SARS-CoV-2 RBD. Position 30 is occupied by an aspartic acid (D) in susceptible humans (*Homo sapiens*), is an asparagine (N) in non-susceptible mice (*Mus musculus*), and an alanine (A) in the avian species (*Aythya fuligula* and *Gallus gallus*). Glutamic acid (E) is present at position 30 in pig (*Sus scrofa*) and Malayan pangolin (*Manis javanicus*), representing intermediate and unknown susceptible species, respectively. Position 34 is conserved as histidine (H) in all susceptible species such as humans, yet has another residue identity in intermediate and non-susceptible species. Species names in orange are susceptible, green are non-susceptible, blue are intermediate susceptibility, and black are unknown.

**Figure 5:**
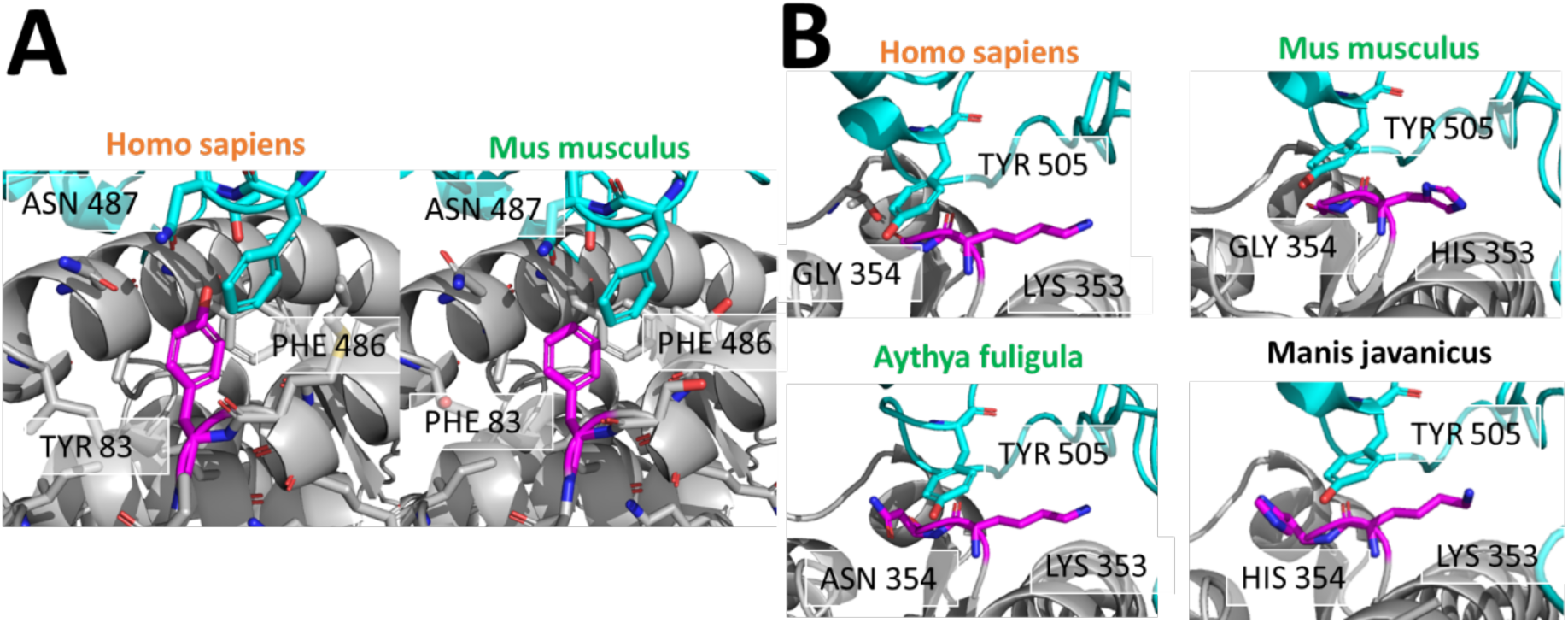
Binding interactions of ACE2 positions 83 and 354 differ across susceptible and non-susceptible species. (**A**) ACE2 position 83 (magenta) is a tyrosine in the human susceptible species (left) and phenylalanine in the non-susceptible mouse species (right). Tyrosine 83 of human ACE2 interacts with asparagine 87 of SARS-CoV-2 RBD, probably via a hydrogen bond. Phenylalanine in mouse ACE2 cannot interact with asparagine 487 due to the lack of a hydrogen bond donor. (**B**) Interactions of tyrosine r505 of the SARS-CoV-2-RBD (cyan) with ACE2 residues 353 and residue 354 (magenta). ACE2 residue 353 is conserved as lysine with the only exception of a histidine in the mouse ACE2. ACE2 residue 354 is a glycine in the susceptible species (human), but an asparagine in non-susceptible duck and chicken, and a histidine in pangolin (unknown susceptibility). Species names in orange are susceptible, green are non-susceptible, and black are unknown.

### Multistate design reveals ACE2 G354 as determinant of susceptibility

It is an evolutionary advantage for SARS-CoV-2 to maintain its ability to infect multiple species. Thus, we hypothesized that the sequence of SARS-CoV-2 RBD is not optimized for a single species but is capable of binding ACE2 of multiple species. Multistate design is a computational approach to test this hypothesis. It allows us to determine the sequence of SARS-CoV-2 RBD that is optimal for binding ACE2 of multiple species. We used Restraint Convergence (RECON) multistate design to test this hypothesis. This method determines how many mutations one protein requires to acquire affinity for multiple targets at once.^25,26^

We adapted this strategy to evaluate the ability of the SARS-CoV-2-RBD to bind non-human ACE2 variants starting from the constraint of the known binding to human ACE2. We hypothesized that engineering a SARS-CoV-2 RBD with binding affinity for ACE2 from non-susceptible species would require more changes to binding interface residues than for susceptible species. To test this hypothesis, we redesigned the SARS-CoV-2 RBD interface sequence using RECON in the presence of the known binder, human ACE2, and ACE2 from other species in turn (**Figure 6A**).

**Figure 6:**
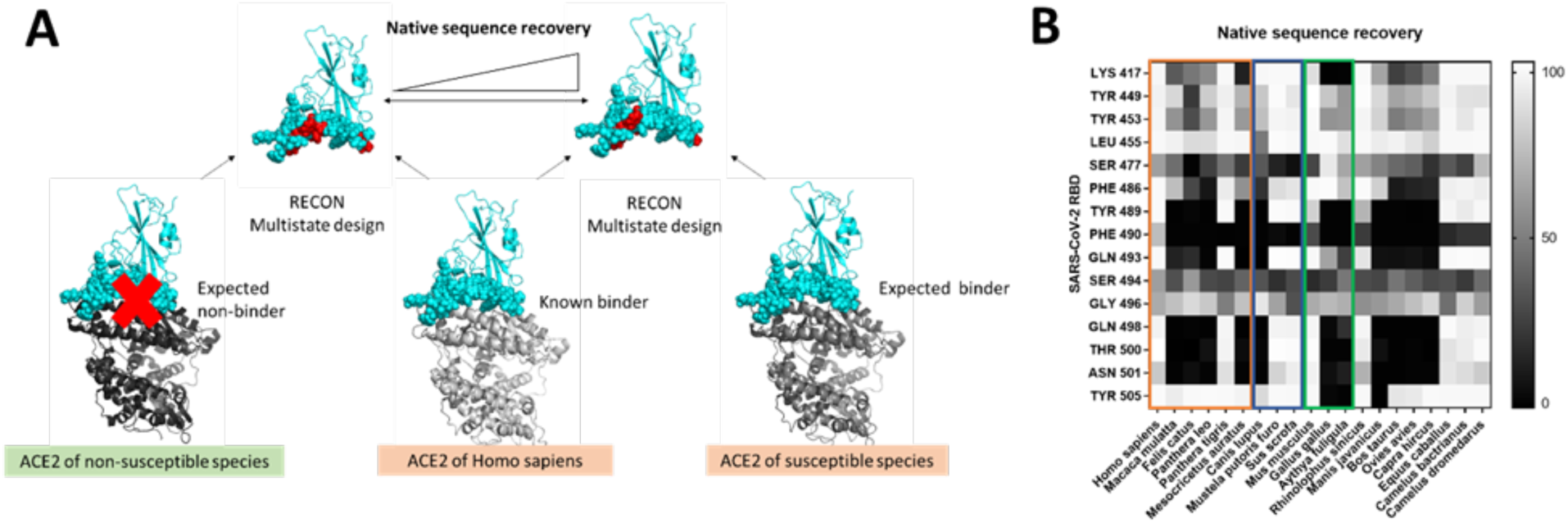
Multistate design reveals SARS-CoV-2 RBD Tyr505 to have low native sequence recovery in non-susceptible duck and chicken. (**A**) RECON multistate design overview. In the presence of ACE2 from two different species the SARS-CoV-2-RBD interface is redesigned. When two true binders are redesigned they should require few sequence changes, thus resulting in a higher native sequence recovery. In contrast, if the native sequence recovery for the interface residues is lower, then many sequence changes are required, indicating that one of the ACE2 proteins is a non-binder. (**B**) Residue-specific native sequence recovery as determined from RECON multistate design against the SARS-CoV-2-RBD complex with human ACE2. Only residues of the SARS-CoV-2-RBD, which are in the protein-protein interface and show changes are depicted. Tyrosine 505 of SARS-CoV-2 RBD shows low native sequence recovery (black) in non-susceptible duck (*Gallus gallus*) and chicken (*Aythya fuligula*). The orange box outlines susceptible species, the blue box outlines species with intermediate susceptibility, and the green box outlines non-susceptible species.

As an initial positive control, the SARS-CoV-2 RBD was redesigned against human ACE2 only. By mutating multiple SARS-CoV-2 RBD residues to improve binding affinity, we tested at each designable position the frequency of native sequence recovery, which measures the fraction of models in which the native SARS-CoV-2 RBD amino acid is retained. This resulted in very few proposed amino acid changes of SARS-CoV-2 RBD to optimally bind human ACE2, indicating that the SARS-CoV-2 RBD sequence overall represents a solution close to optimal (**Figure 6B**). The exception is valine 503, for which more polar amino acids were deemed optimal. This valine, however, is near a glycosylation site at asparagine 322 in ACE2 at the SARS-CoV-2 and ACE2 interface (**Extended Data Figure 5**). Since glycans are not incorporated into the RECON multistate design technique, this valine 503 may have a higher affinity binding partner when considering the presence of ACE2 glycosylation sites..

Designing SARS-CoV-2 RBD in the presence of ACE2 from additional species revealed that ACE2 from a number of species have lower sequence recovery (including non-susceptible species such as duck and chicken, but also hamster, macaque, cat, lion and dog). When evaluating residue-specific interactions based on the native sequence recovery from RECON multistate design, tyrosine 505 shows no sequence recovery in avian species as compared to the human ACE2 control. This tyrosine interacts very prominently with lysine 353 in ACE2, however this residue is highly conserved across all species examined (**Table 1**). Tyrosine 505 also interacts less strongly with glycine 354, which is occupied by an asparagine in the avian species (chicken and duck) (**Table 1 and Figure 5B**). This secondary interaction might explain the differences in native sequence recovery. However, another experimentally verified non-susceptible species, the mouse (*Mus musculus*), has a high degree of sequence recovery, similar to human ACE2. This suggests that other factors beyond residue-residue interactions of ACE2 and SARS-CoV-2 RBD at the interface may determine susceptibility to infection, at least in the mouse, and that differences in RECON multistate design explain only partially differences in species susceptibility to SARS-CoV-2 infection.

### ACE2 glycosylation at N90 and N322 as determinants of susceptibility

As a final additional approach to structurally evaluate differences in species susceptibility, we investigated the predicted glycosylation profiles of various species in comparison to human ACE2. Protein glycosylation is increasingly recognized as a critical contributor to receptor-ligand interactions;^27^ however, given the challenges in identifying glycans in protein crystal structures, glycosylation has received considerably less attention than SARS-CoV-2 RBD and ACE2 protein-protein interactions. Naturally occurring glycans as posttranslational modifications are not fully visible in crystal structures. Normally only the first *N*-actylglucosamine is visible or no sugar moiety can be observed, or glycosylation sites are mutated prior to crystallization. In the crystal structures of the human ACE2 used here, a sugar moiety bound to an asparagine at a surface exposed NXT/S sequon was seen three times in proximity to the binding interface on the ACE2. To understand whether the ACE2 of other species have similar glycosylation patterns, glycosylation was predicted using NetNGlyc 1.0, a neural network for predicting N-glycosylation sites, and compared to the glycosylation patterns of human ACE2.^28^ Residues 53, 90, 103, and 322 were identified as glycosylation sites in human ACE2, with 53, 90, and 322 demonstrating glycosylation in the crystal structure (PDB: 6M0J and 6LZG)^14^ (**Table 2**). Other susceptible species were quite similar to this pattern, except for position 103, which is only predicted to be glycosylated in humans and rhesus macaques. Among known susceptible species, only Golden Syrian hamster ACE2 lacks predicted glycosylation in position 322. At position 90, all susceptible species were predicted to be glycosylated and all non-susceptible and intermediate susceptibility species were non-glycosylated. Interestingly, ACE2 from the non-susceptible mouse, despite not showing significant differences in predicted binding energy or RECON multistate analysis compared to susceptible species, is predicted to lack glycosylation at residues 90 and 322, distinguishing it from ACE2 of nearly all susceptible species. This suggests a potential mechanism by which mice may be non-susceptible despite having similar binding energy and SARS-CoV-2 native sequence recovery to susceptible species.

**Table 2:**
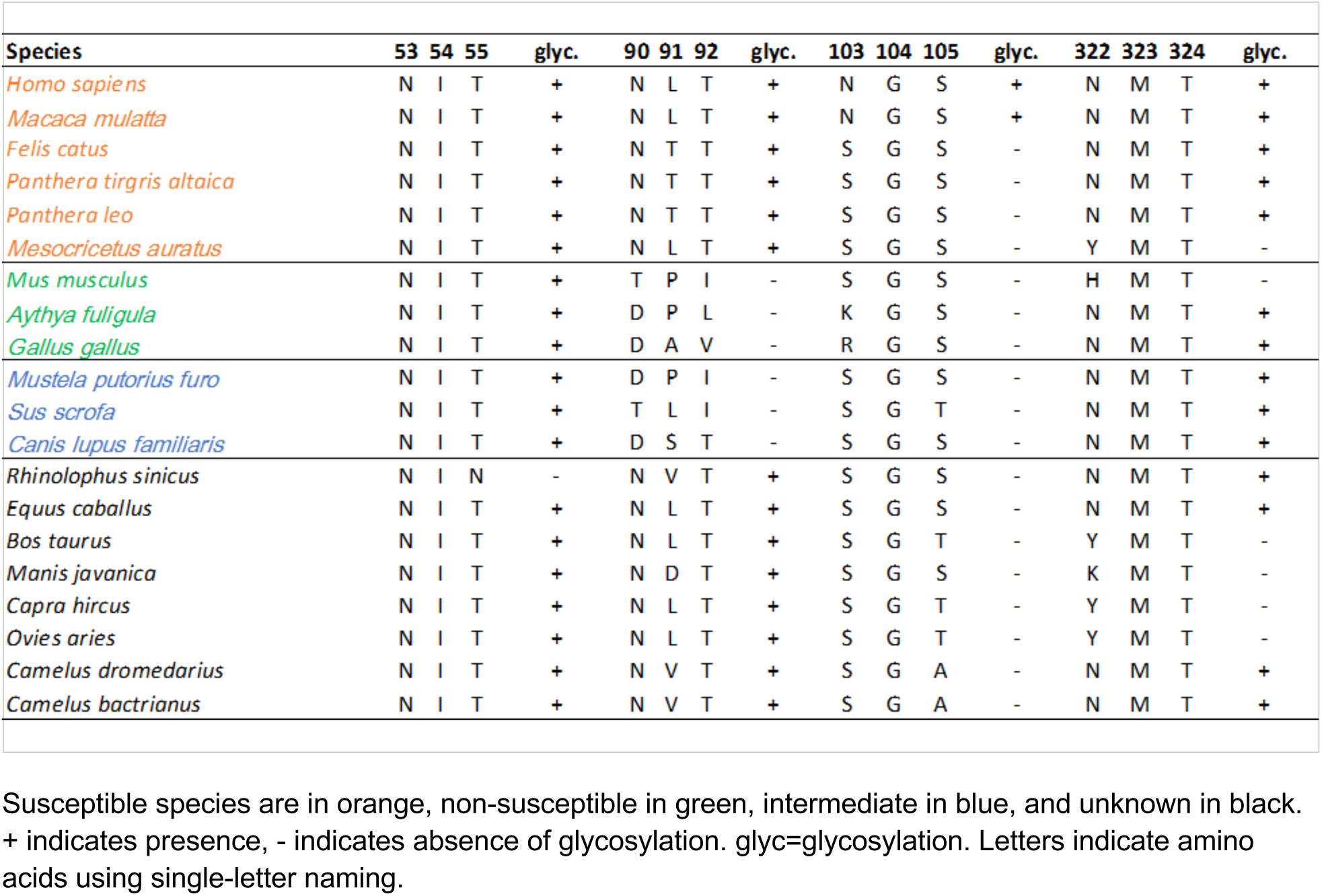
Predicted glycosylation profiles for ACE2 amino acid positions 53, 90, 103 and 322.

### A SARS-CoV-2 susceptibility score predicts species at risk

Taken together, results of these studies reveal a set of key ACE2 residues important for interaction with SARS-CoV-2 RBD and for which differences help discriminate susceptible from non-susceptible species. These differences include ACE2 amino acid positions 30 and 83, which exhibit differential residue-residue binding energy, position 354, which exhibits low native sequence recovery in interaction with SARS-CoV-2, and positions 90 and 322, which exhibit differences in glycosylation. Using these key residues in aggregate, we developed a SARS-CoV-2 susceptibility score based on similarity to the human ACE2 sequence using the BLOSUM62 similarity matrix (**Table 3**).^29^ This analysis revealed that experimentally validated non-susceptible species have in fact the lowest susceptibility scores, while species with previously demonstrated intermediate susceptibility have intermediate susceptibility scores. Using the lowest score of the susceptible species, 23, as the lower cutoff for susceptibility and the highest score of non-susceptible species, 11, as the upper cutoff for non-susceptibility, we extended these results to species with unknown susceptibility. This revealed high scores in the susceptible range for the Chinese horseshoe bat (*Rhinolophus sinicus*), horse (*Equus caballus*), and camels (*Camelus dromedarius* and *Camelus bactrianus*) and intermediate susceptibility scores for the Malayan pangolin (*Manis javanica*), cow (*Bos taurus*), goat (*Capra hircus*), and sheep (*Ovis aries*).

**Table 3:** Key residues of aligned ACE2 proteins with calculated SARS-CoV-2 susceptibility score for each species.

|  | 30 | 83 | 90 | 322 | 354 | **Susceptibility score** |
| --- | --- | --- | --- | --- | --- | --- |
| *Homo sapiens* (Human) | D | Y | N | N | G | - |
| *Macaca mulatta* (Rhesus Macaque) | D | Y | N | N | G | 31 |
| *Felis catus* (House cat) | E | Y | N | N | G | 27 |
| *Panthera tigris altaica* (Tiger) | E | Y | N | N | G | 27 |
| *Panthera leo* (Lion) | E | Y | N | N | G | 27 |
| *Mesocricetus auratus* (Golden Syrian Hamster) | D | Y | N | Y | G | 23 |
| *Mus musculus* (Mouse) | N | F | T | H | G | 11 |
| *Aythya fuligula* (Duck) | A | F | D | N | N | 8 |
| *Gallus gallus* (Chicken) | A | F | D | N | N | 8 |
| *Mustela putorius furo* (Ferret) | E | Y | D | N | R | 14 |
| *Sus scrofa* (Pig) | E | Y | T | N | G | 21 |
| *Canis lupus familiaris* (Dog) | E | Y | D | N | G | 22 |
| *Rhinolophus sinicus* (Chinese horseshoe bat) | D | Y | N | N | G | 31 |
| *Equus caballus* (Horse) | E | Y | N | N | G | 27 |
| *Bos taurus* (Cow) | E | Y | N | Y | G | 19 |
| *Manis javanica* (Malayan pangolin) | E | Y | N | K | H | 13 |
| *Capra hircus* (Goat) | E | Y | N | Y | G | 19 |
| *Ovis aries* (Sheep) | E | Y | N | Y | G | 19 |
| *Camelus dromedarius* (Arabian Camel) | E | Y | N | N | G | 27 |
| *Camelus bactrianus* (Bactrian Camel) | E | Y | N | N | G | 27 |

Susceptible (orange), non-susceptible (green), intermediate (blue), and unknown (black/grey) species are indicated.

To permit wider use of this susceptibility score for evaluation of additional species with unknown susceptibility, including those species that in the future may be of particular concern, we developed an implementation of the susceptibility score algorithm in R for public use. This implementation takes as input human ACE2 aligned with ACE2 of another species of interest and provides a susceptibility score using differences in ACE2 positions 30, 83, 90, 322, and 354. R code for implementation of this algorithm as a graphical user interface is available in Supplemental Methods.

## DISCUSSION

Here we tested the hypothesis that differences in ACE2 proteins across various species alter structural interactions with SARS-CoV-2 RBD, leading to differences in species susceptibility to SARS-CoV-2 infection. Our results, combining prior knowledge of experimentally validated differences in species susceptibility with multiple methods of determining effects on ACE2 structure and interaction with SARS-CoV-2 RBD, reveal five key residues that in aggregate help discriminate susceptibility across species. These include ACE2 positions 30, 83, and 354, which exhibit alterations in binding energy, and positions 90 and 322, which exhibit alterations in glycosylation that likely contribute to differences in interactions at the interface. Taken together, our results provide insight into the molecular determinants of species susceptibility to SARS-CoV-2 infection and have important implications for identification of key residues for therapeutic targeting and determining susceptibility of additional species to infection.

Our study has several unique features that permit rigorous evaluation of differences in species susceptibility to infection. Prior studies have similarly performed ACE2 sequence alignments across species and modeled structural effects of the amino acid changes on the SARS-CoV-2 and ACE2 interface.^7,30-35^ However, our study integrates experimentally validated susceptibility to SARS-CoV-2 with in-depth structural analyses to determine critical ACE2 residues for infection. In addition, we performed multiple structural analyses, including residue-residue interactions, RECON multistate design, and glycosylation analysis, to rigorously determine the structural basis for species differences in ACE2 interaction with SARS-CoV-2 RBD. Prior studies of ACE2 sequence alignment with limited structural modeling have suggested that pigs are susceptible to infection,^36^ and that hamsters and house cats are in an intermediate risk group.^37^ Recent experimental work with direct inoculation, however, has demonstrated that pigs are non-susceptible,^5^ and that house cats and Golden Syrian hamsters are susceptible.^5,7^ We identified key residues on which to build a susceptibility score that closely matches experimentally verified *in vivo* susceptibility, including predicting an intermediate susceptibility of the pig and higher susceptibility of house cats and Golden Syrian hamsters.

A key principle revealed by our findings is the importance of using multiple methods for determining the structural basis for differences in ACE2 interaction with SARS-CoV-2 RBD. For example, although calculated binding energy, protein stability, and RECON multistate design of SARS-CoV-2 RBD in complex with duck and chicken ACE2 distinguished non-susceptible chicken and duck ACE2 from susceptible species, mouse ACE2 did not fit the pattern of other non-susceptible species. However, analysis of ACE2 protein glycosylation revealed two residues, 90 and 322, for which differences in mouse ACE2 distinguished it from susceptible species. In addition, combining ACE2 sequence alignment, GroupSim calculations, and residue-residue interaction modeling identified residues 30 and 83, which are distinctly different in all non-susceptible compared to susceptible species. Differences in these residues in non-susceptible species result in decreased binding energy with SARS-CoV-2 RBD. Although no single residue appears capable of explaining the difference in susceptibility to SARS-CoV-2 infection across species, in combination amino acid positions 30, 83, 90, 322, and 354 can help distinguish susceptible from non-susceptible species, as reflected by the calculated susceptibility score, which was lower in non-susceptible species and intermediate in those species with intermediate susceptibility.

Our findings have important implications for determining infectability of animals with heretofore unknown susceptibility to SARS-CoV-2 infection. Determining such susceptibility is critical to prevent disruption to food supplies, identify optimal animal models for research, aid in the search for intermediate hosts, and enhance identification of potential animal reservoirs that can propagate transmission.^38^ We applied our infection susceptibility score to several important species with unknown susceptibility to date. These data suggest that cows (*Bos taurus*), Malayan pangolin (*Manis javanica*), and goats (*Capra hircus*) have intermediate susceptibility to infection, while Chinese horseshoe bats (*Rhinolophus sinicus*), horses (*Equus caballus*), and camels (*Camelus dromedarius* and *Camelus bactrianus*) have higher susceptibility. Although the ultimate test is direct exposure of live animals to evaluate infectability and transmissability,^5,7^ this is complicated by the need for BSL3 containment and is quite costly and challenging with larger animals. Observational studies and case reports could also help provide evidence of susceptibility. Indeed, our results suggest that horses and camels should be tested and/or closely monitored for evidence of Covid-19 infection. The close interaction of these animals with humans, and the importance of these animals as domestic companions and laborers worldwide make determination of their susceptibility an urgent need. The use of the susceptibility score developed here can also be applied to additional species of interest to help direct resources for focused research and protection efforts in the future.

ACE2 residues identified in this paper that provide a structural basis to differences in species susceptibility to infection reveal important insights into the SARS-CoV-2 RBD and ACE2 structural interaction and potential for therapeutic targeting. By incorporating differences in species susceptibility into the structural analysis, our findings enhance the potential to identify particularly important residues mediating the ACE2 and SARS-CoV-2 RBD interaction. Indeed, although GroupSim scores were not used in the structural analysis, three of the five key identified residues (30, 83, and 90) from the structural modeling are in the top scoring ACE2 positions by GroupSim score. This suggests that the amino acids at these positions in ACE2 differ significantly between susceptible and non-susceptible species, consistent with an important contribution of these residues to differences in susceptibility. Amino acid positions 30 and 83 of ACE2 in particular exhibited large differences in residue-residue interaction binding energies between susceptible and non-susceptible species. Asp30 on ACE2 interacts with residues Lys417, Phe456, and Tyr473 of SARS-CoV-2 RBD, and ACE2 Tyr83 interacts with Asn487 and Tyr489 of SARS-CoV-2 RBD. These amino acids mark sites of SARS-CoV-2 interaction with ACE2 that may be important for development of antibody-based therapies or small molecule inhibitors.

Applying a multistate design algorithm to probe the SARS-CoV-2-RBD interactions for their ability to cross-bind to ACE2 of multiple species yielded several novel observations. First, this technique identified ACE2 position 354 as an important site for differentiating binding and non-binding ACE2 of different species to SARS-CoV-2 RBD. Second, this approach demonstrated that the SARS-CoV-2 RBD sequence is nearly optimal for binding to human ACE2 compared to other species. This is a remarkable finding, and likely underlies the high transmissibility of this virus amongst humans. This finding is also consistent with recent results that compared SARS-CoV and SARS-CoV-2 and determined that a number of differences in the SARS-CoV-2 RBD have made it a much more potent binder to human ACE2 through the introduction of numerous hydrogen bonding and hydrophobic networks.^39^

Although ACE2 and SARS-CoV-2 RBD interactions are critical to SARS-CoV-2 infection,^9,10,16^ differences in other factors across species may also contribute to differences in susceptibility. This includes differences in ACE2 expression levels^40^ and differences in the protein sequence of TMPRSS2, a protein that contributes to viral and host cell membrane fusion through cleavage of spike protein.^15,41^ With further experimental and observational data on infectability of currently unknown species, the susceptibility score we have developed can also help determine species for which differences in ACE2 protein may not inadequately predict differences in susceptibility. For these species future studies could compare differences in expression levels of ACE2 and/or differences in TMPRSS2 structure. These structural comparisons of TMPRSS2, however, will require elucidation of the protein crystal structure, which is not yet available.

## CONCLUSION

We combined in-depth structural analyses with knowledge of varying species susceptibility to SARS-CoV-2 infection to determine key structural determinants of infection susceptibility. First, we identified multiple key residues mediating structural interactions between ACE2 and SARS-CoV-2 RBD. Differences in these residues were used to generate a susceptibility score that can help predict animals with elevated risk of infection for which we do not yet have experimental evidence of susceptibility, including horses and camels. Finally, we have demonstrated that SARS-CoV-2 is nearly optimal for binding ACE2 of humans compared to other animals, which may underlie the highly contagious transmissibility of this virus amongst humans. Taken together, results of these studies identify key structural regions of the ACE2 and SARS-CoV-2 interaction for therapeutic targeting and for identifying animal species on which to focus additional research and protection efforts for environmental and public health.

## METHODS

### ACE2 protein alignment

Protein sequence accession numbers and corresponding FASTA files from multiple species (**Extended Data Table 2**) were pulled from NCBI using Batch Entrez. In the absence of a published sequence and accession number, ACE2 protein sequence for the lion (*Panthera leo*) was assembled using TBLASTN (National Center for Biotechnology Information) with tiger ACE2 protein sequence as the query (**Extended Data Table 3**). Protein sequences were loaded into EMBL-EBI web interface implementation of MAFFT for multiple sequence alignment using default settings (https://www.ebi.ac.uk/Tools/msa/mafft/).^20^ Resulting alignment was uploaded to ESPript 3.0 to generate a graphical version of the alignment (http://espript.ibcp.fr/ESPript/ESPript/), including annotation of secondary structure based on Protein Data Bank (PDB) structure 1r42 of human ACE2.^42^ A treedyn format tree diagram representing similarity of ACE2 protein sequence across species was generated using phylogeny.fr (https://www.phylogeny.fr/).^43,44^ NCBI Taxonomy Browser was used to generate a taxonomic tree of phylogenetic relationships amongst species as a Phylogeny Inference Package (PHYLIP) tree.^45^ Final visualization was performed using the interactive Tree of Life (iTOL) tree viewer v 5.5.1 (https://itol.embl.de/).^46^

### Quantification of amino acid differences in alignment of susceptible and non-susceptible species

Quantification of amino acid positions in the ACE2 protein alignment that optimally distinguish susceptible versus non-susceptible species was performed using GroupSim.^21^ Values from 0 to 1 were obtained with 1 assigned to the position that best stratifies susceptible and non-susceptible species. Values are weighted by the BLOSUM62 similarity matrix to incorporate similarity of amino acids properties.^29^

### Homology modeling of ACE2-SARS-CoV2 co-crystal structures using RosettaCM

ACE2 of human and non-human species was modeled based on two co-crystal structures of SARS-CoV-2-RBD with the human ACE2 (PDB-IDs 6LZG and 6M0J).^14^ One co-crystal structure (PDB-ID 6VW1) was excluded due to its lower resolution as compared to the aforementioned structures. The target sequences were threaded over the ACE2-SARS-CoV-2-RBD co-crystal structure, which was first relaxed with backbone constraints using RosettaRelax.^47^ A total of 1000 homology models were constructed using RosettaCM, and subsequently relaxed with backbone constraints.^47,48^ Of these, 25 models were selected based on the total energy as a measure of protein stability, predicted binding energy, and Cα-root mean square deviation (Cα-RMSD) to the best scoring model (**Extended Data Figure 3**). The SARS-CoV-2-RBD-ACE2 complex was optimized using a rigid-body docking with limited degrees for rotational and torsional sampling.^49,50^ A final ensemble of 100 models was selected based on the total energy as measure of protein stability, predicted binding energy and Cα-RMSD to the best scoring model (**Extended Data Figure 4**). The pairwise binding interaction between SARS-CoV-2 and ACE2 was evaluated by retrieving the decomposed Rosetta scores for each residue. The protocol was tested by modeling the human ACE2 in complex with SARS-CoV-2-RBD, and evaluating the recovery of predicted binding energy, total energy, and residue-residue interactions in the interface.

### Calculation of sequence recovery from Restraint Convergence (RECON) multistate design

RECON multistate design was carried out as reported previously for each susceptible, non-susceptible, intermediate, and unknown species against the human SARS-CoV-2-RBD-ACE2 complex.^25,26,51^ As a control, this was also performed solely using the human SARS-CoV-2-RBD-ACE2 complex. A total of 5000 models were sampled and trajectories with final models that scored lower than -2400 REU were evaluated. The native sequence recovery was calculated for each pairwise experiment and also for the control run for the SARS-CoV-2-RBD complex with the human ACE2 (**Extended Data Figure 6**).

All protocols were executed using Rosetta-3.12 (www.rosettacommons.org). Evaluation was performed using the numpy, pandas, matplotlib and seaborn libraries in Python 3.7, PyMOL 2.7^52-54^ and GraphPad Prism version 8.3.0 for Windows (GraphPad Software, San Diego, California). Example commands and RosettaScripts protocols can be found in the Supplementary Methods.

### Prediction of glycosylation sites

The NetNGlyc 1.0 server (http://www.cbs.dtu.dk/services/NetNGlyc/) was used to predict glycosylation sites.^28^ Based from the observation that asparagine in positions 53, 90, and 322 carried glycosylation in the crystal structures PDB: 6LZG and 6M0J, and scored with high confidence from NetNGlyc 1.0, these were selected as reliably glycosylated. Position 103 was included, as it was strongly predicted to be glycosylated by NetNGlyc 1.0, although no glycosylation was observed in the crystal structures. Furthermore, it was evaluated whether the NxT/S sequons were surface accessible and in proximity to the ACE2-SARs-CoV-2-RBD binding interface.

### SARS-Cov-2 susceptibility score calculation

Using identified ACE2 key amino acid positions 30, 83, 90, 322, and 354 in the alignment of ACE2 across species, a global susceptibility score was calculated as the sum of the Blosum62 scoring matrix substitutions for the amino acid at each position compared to the human ACE2 sequence.^29^ This was calculated for each species, with higher scores suggesting greater susceptibility. An R implementation of this susceptibility score algorithm was also developed in RStudio. The software takes as input alignment of human ACE2 protein sequence with ACE2 of another species of interest and provides a susceptibility score as output. Susceptibility scores of species examined in this manuscript are also graphically demonstrated as reference. Code for implementing this algorithm in R as a graphical user interface is available in Supplemental Methods.

### Statistical analysis

Contingency testing was performed with Fisher’s exact test as a two-sided comparison and alpha equal to 0.05 using GraphPad Prism version 8.2.1 (GraphPad Software, Inc.).

## Acknowledgements

The authors would like to thank Erkan Karakas for useful suggestions at the beginning of the project and Melissa Farrow for helpful suggestions in the conceptualization of the study. This work was supported in part by the National Institutes of Health under awards F32HL144048-01 (MRA), DK117147 (WC), UH3TR002097 and U01TR002383 (JPW and JAB), U19AI117905, U01AI150739, and R01AI141661 (JM and CTS), R35GM127087 (CM and JAC), and DP2HL137166 (MSM). The work was also supported by the American Heart Association 20PRE35080177 (CDS) and EIA34480023 (MSM). The views expressed are solely those of the authors.

## Author Contributions

JAB and JPW conceptualized the study; MRA, CTS, WC, MSM, JM, JPW, JAB, CM, and JAC made substantial contributions to experimental design; MRA and CTS conducted the majority of the experiments with help from WC, JM, CDS, JAC, JAB, and JPW; MRA, CTS, JAC, WC, CM, and JM significantly contributed to the data acquisition and interpretation; MRA and CTS drafted the manuscript with WC, MSM, JM, JPW, and JAC contributing critical revisions. All authors approved the final version.

## Data and Code Availability

The datasets generated and/or analyzed during the current study are available from the corresponding author on reasonable request. R code for the susceptibility score algorithm is available in Supplemental Methods.

## Additional Information

Supplementary Information is available for this paper.

## Extended Data Tables

**Extended Data Table 1:** GroupSim scores of aligned ACE2 amino acid positions with greatest differences between susceptible and non-susceptible species.

| Amino Acid | GroupSim score |
| --- | --- |
| 682 | 1.000 |
| **79** | 0.661 |
| 568 | 0.643 |
| 337 | 0.643 |
| 286 | 0.643 |
| 246 | 0.554 |
| **34** | 0.548 |
| 92 | 0.536 |
| 751 | 0.518 |
| 593 | 0.494 |
| 641 | 0.464 |
| **83** | 0.464 |
| 536 | 0.458 |
| 709 | 0.446 |
| 321 | 0.446 |
| 637 | 0.429 |
| 90 | 0.411 |
| **24** | 0.406 |
| 689 | 0.405 |
| 91 | 0.404 |
| 253 | 0.393 |
| 139 | 0.393 |
| 782 | 0.381 |
| 728 | 0.381 |
| 113 | 0.381 |
| 228 | 0.380 |
| 752 | 0.375 |
| 59 | 0.375 |
| **31** | 0.375 |
| 653 | 0.371 |
| **30** | 0.361 |
| 765 | 0.357 |
| 675 | 0.357 |
| **329** | 0.357 |
| 214 | 0.357 |

Bold amino acids are structurally predicted or experimentally validated to mediate ACE2 interactions with SARS-CoV-2 spike protein. Amino acid numbers correspond to *Homo sapiens* ACE2.

**Extended Data Table 2:** Species and accession numbers used for ACE2 protein sequence alignment.

| <u>Common name</u> | <u>Genus species</u> | <u>Accession number</u> |
| --- | --- | --- |
| Human | *Homo sapiens* | NP_001358344.1 |
| House cat | *Felis catus* | XP_023104564.1 |
| Tiger | *Panthera tigris altaica* | XP_007090142.1 |
| Lion | *Panthera leo* | No accession number |
| Golden Syrian hamster | *Mesocricetus auratus* | XP_005074266.1 |
| Rhesus macaque | *Macaca mulatta* | NP_001129168.1 |
| Mouse | *Mus musculus* | NP_081562.2 |
| Duck | *Aythya fuligula* | XP_032058386.1 |
| Chicken | *Gallus gallus* | XP_416822.2 |
| Ferret | *Mustela putorius furo* | NP_001297119.1 |
| Pig | *Sus scrofa* | NP_001116542.1 |
| Dog | *Canis lupus familiaris* | NP_001158732.1 |
| Chinese horseshoe bat | *Rhinolophus sinicus* | AGZ48803.1 |
| Horse | *Equus caballus* | XP_001490241.1 |
| Cow | *Bos taurus* | XP_005228485.1 |
| Malayan pangolin | *Manis javanica* | XP_017505746.1 |
| Goat | *Capra hircus* | NP_001277036.1 |
| Sheep | *Ovis aries* | XP_011961657.1 |
| Arabian camel | *Camelus dromedarius* | XP_010991717.1 |
| Bactrian camel | *Camelus bactrianus* | XP_010966303.1 |

Susceptible (red), non-susceptible (green), intermediate (blue), and unknown (black) species are indicated.

**Extended Data Table 3:**
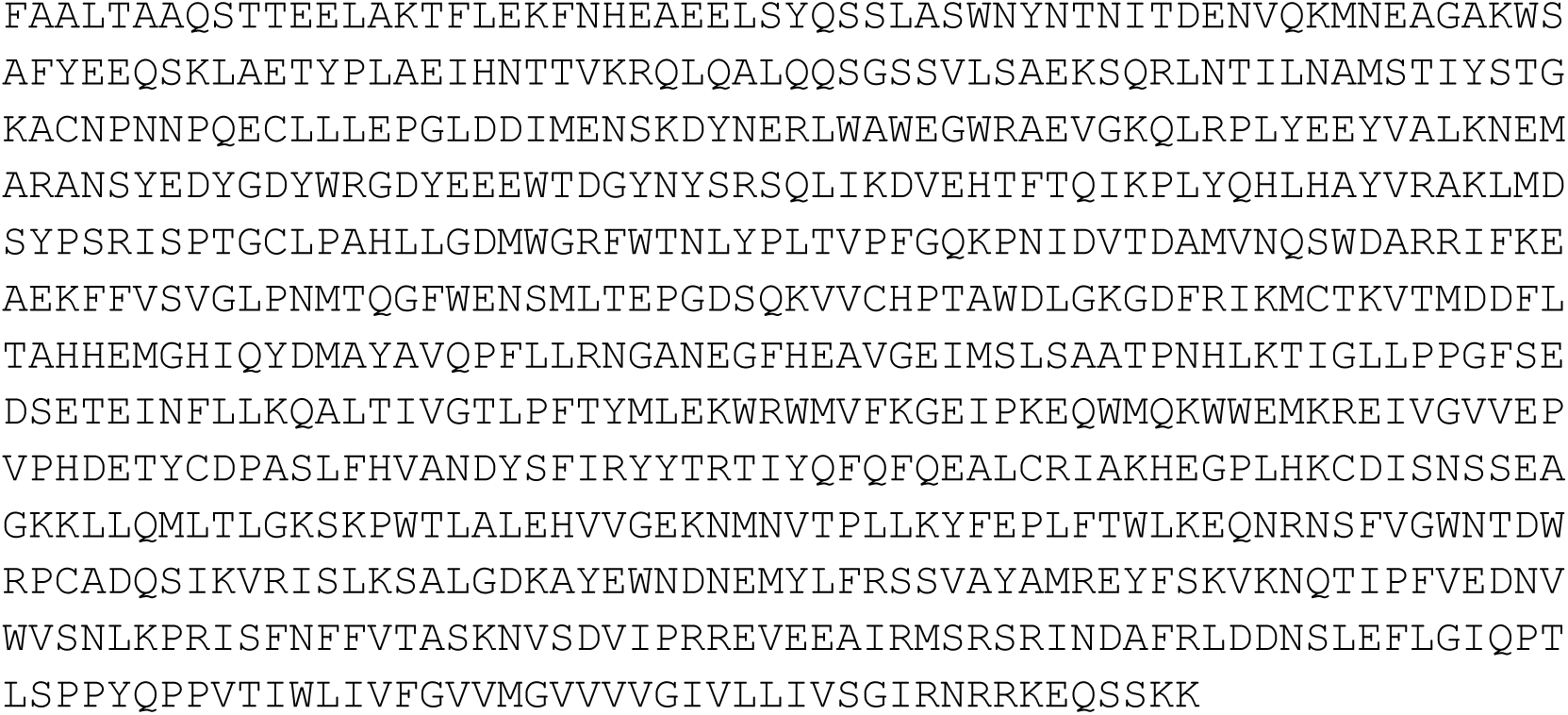
*Panthera leo* ACE2 protein sequence.

## Extended Data Figures

**Extended Data Figure 1:**
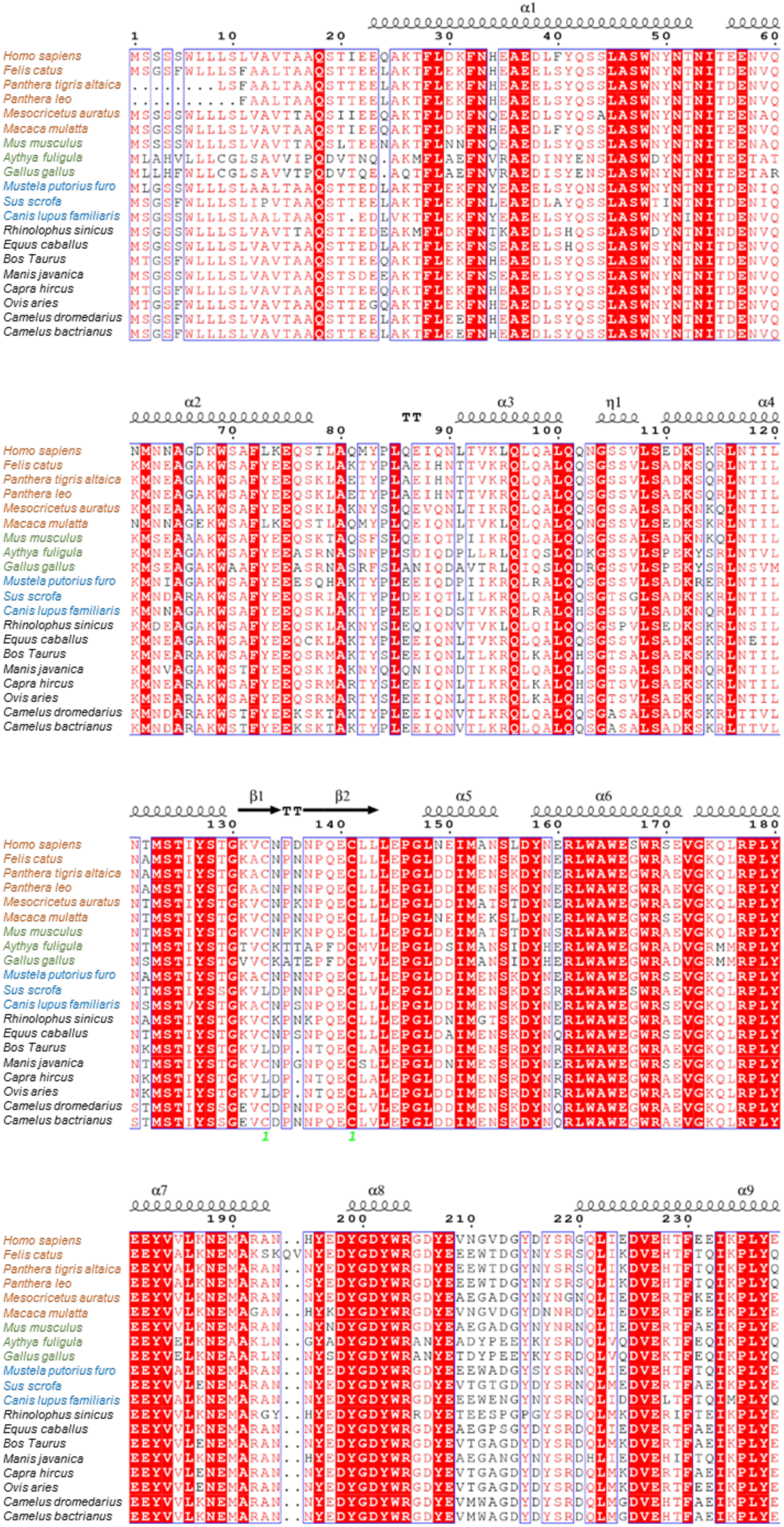
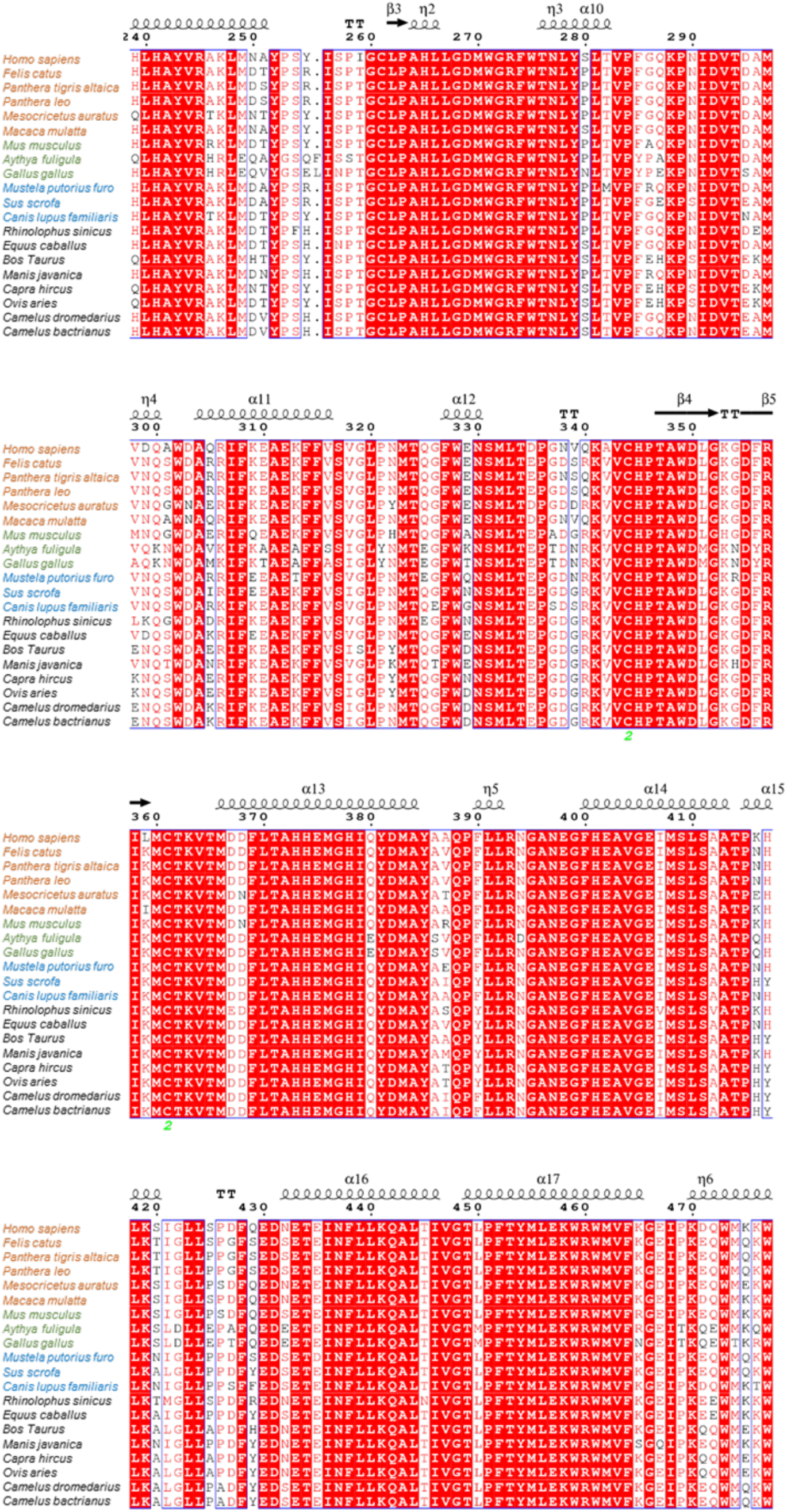
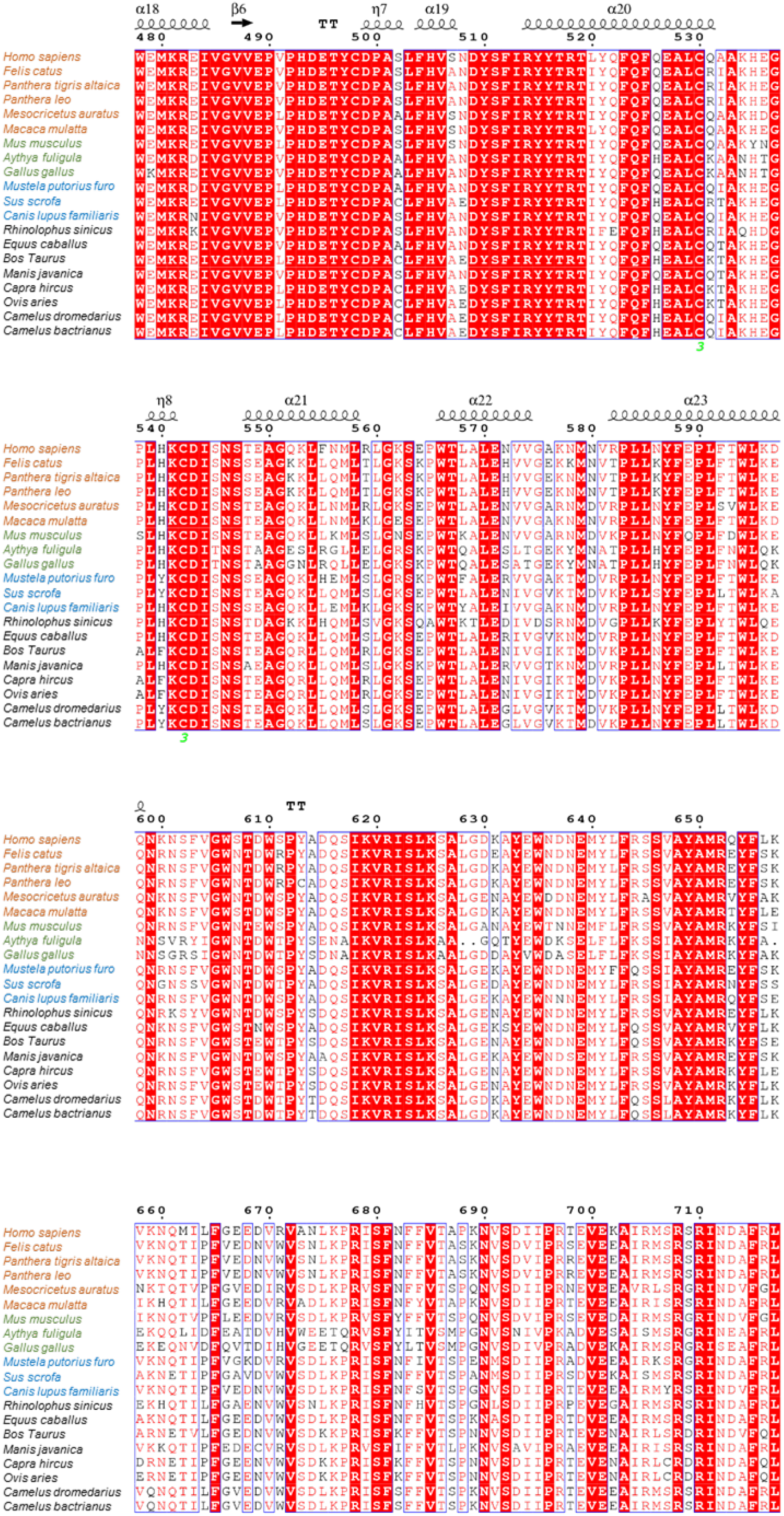
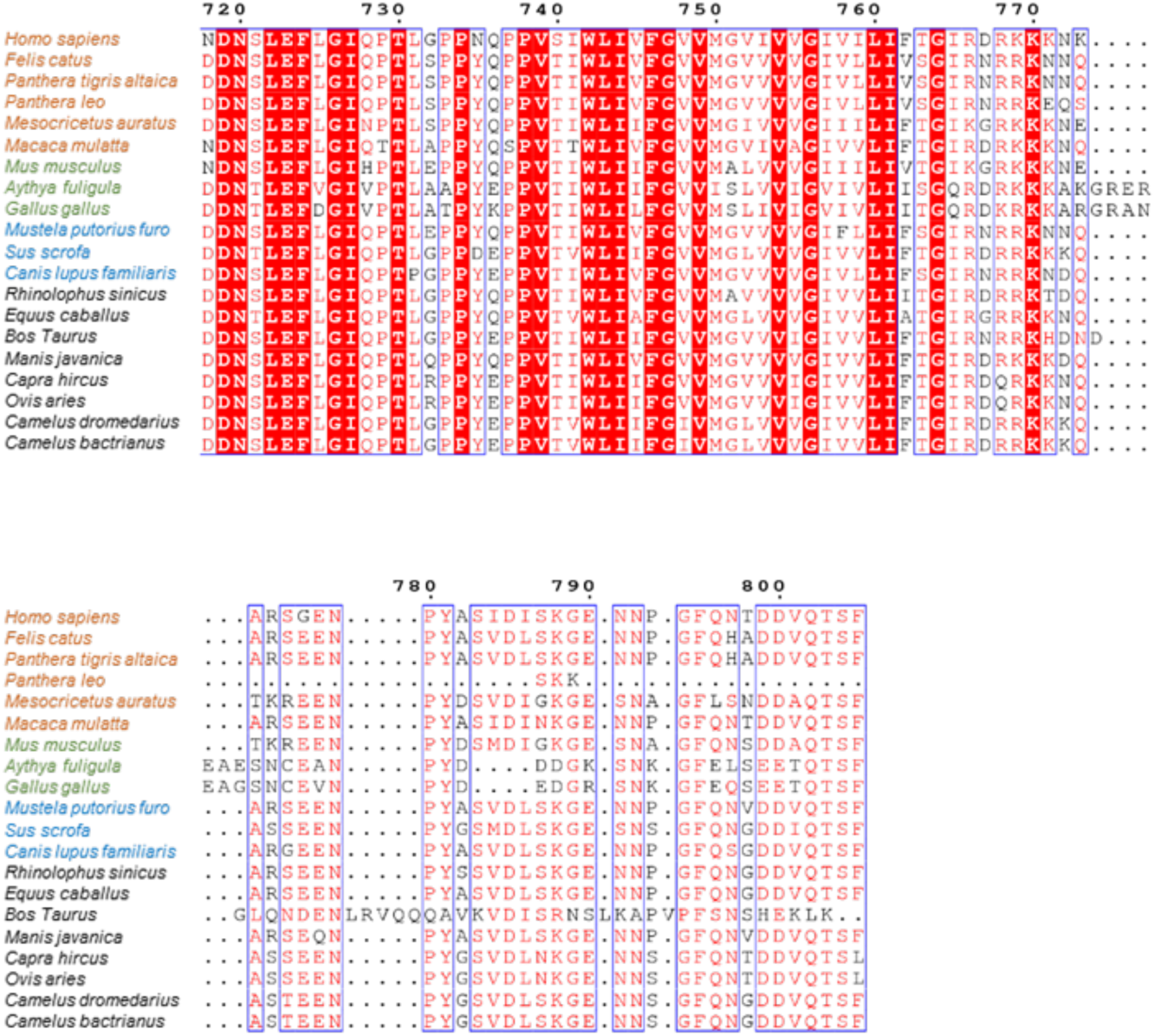
ACE2 protein sequence alignment from susceptible (orange), non-susceptible (green), intermediate susceptible (blue), and unknown susceptible (black) species. MAFFT alignment with visualization using ESPript. Secondary structure elements defined based on human ACE2 (PDB 1r42). Spirals represent α or 310 helices and arrows represent beta strands.

**Extended Data Figure 2:**
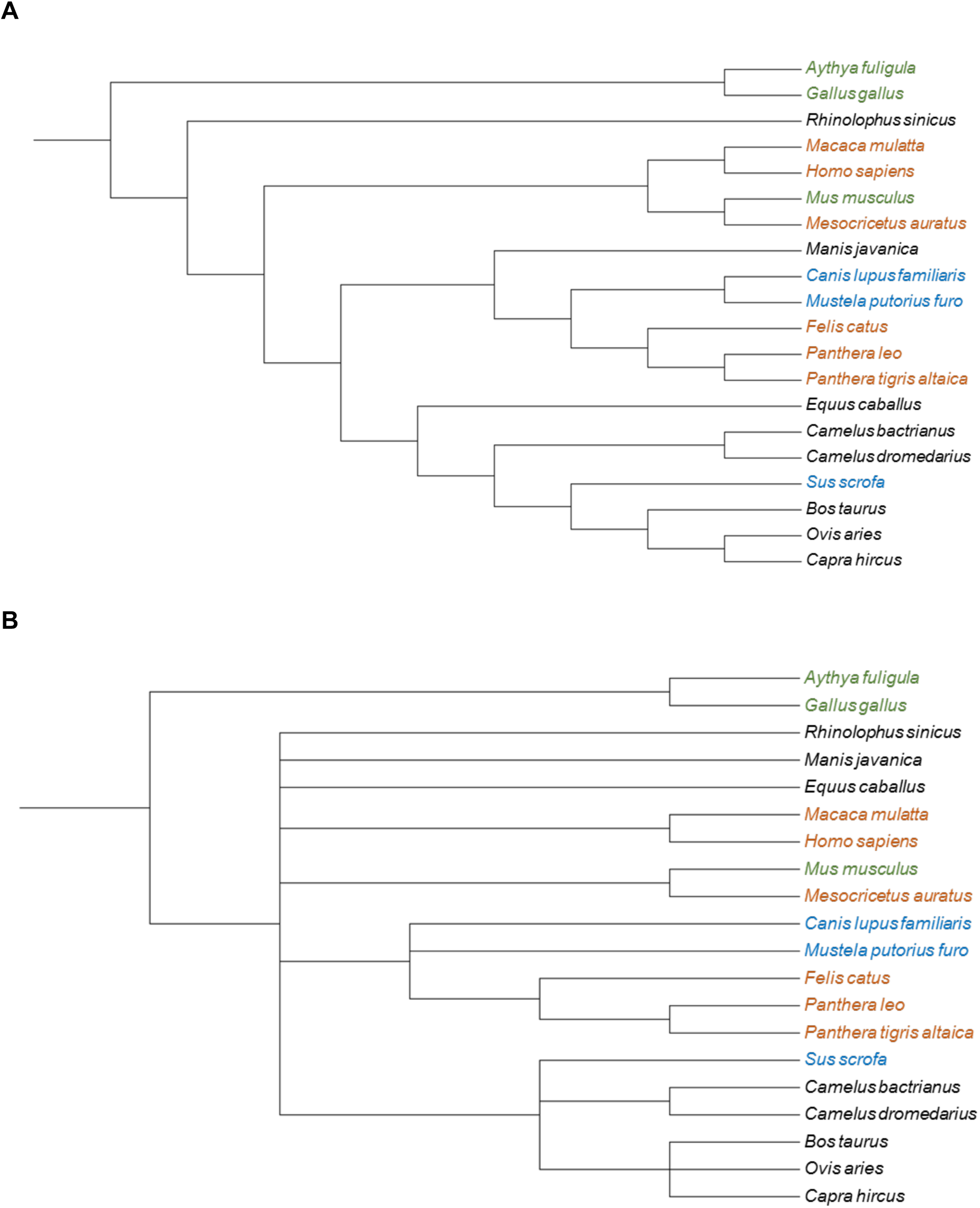
Differences in ACE2 protein sequences and phylogenetic relationships are similar across species. (A) Dendrogram of ACE2 protein sequence comparisons and (B) phylogenetic relationships of susceptible (orange), non-susceptible (green), intermediate susceptibility (blue), and unknown susceptibility (black) to SARS-CoV-2 infection.

**Extended Data Figure 3:**
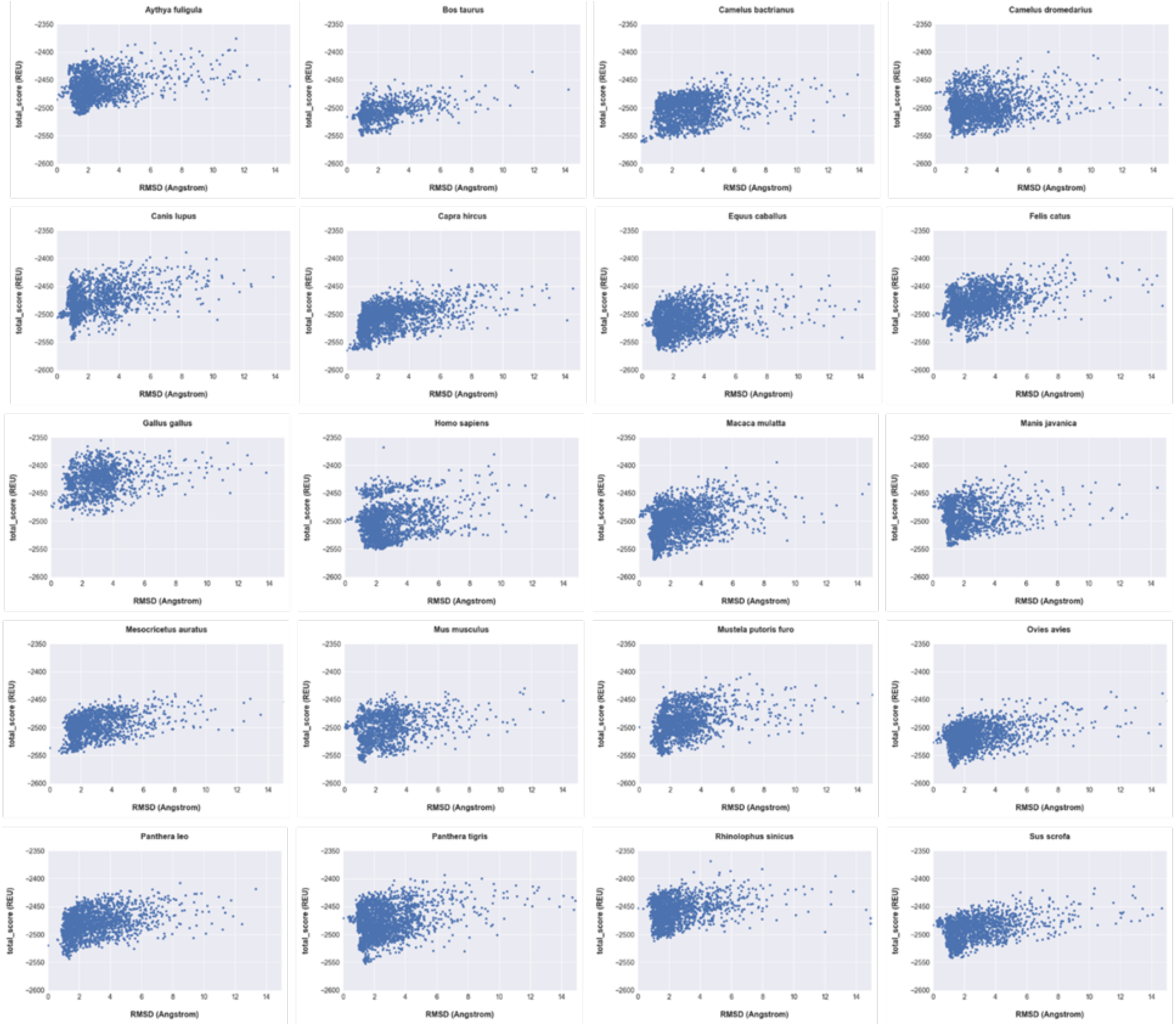
Construction of SARS-CoV-2-RBD-ACE2 complex models for each species resulted in comparable high-quality models. Models were evaluated for their Cα-root mean square deviation (Cα-RMSD) as a measure of similarity to the best performing model by predicted binding energy versus calculated protein stability. Models in the lower left quadrant of the plots show good convergence of calculated protein stability and similarity, and were thus selected for re-docking of SARS-CoV-2-RBD to the respective ACE2 as in Extended Data Figure 4.

**Extended Data Figure 4:**
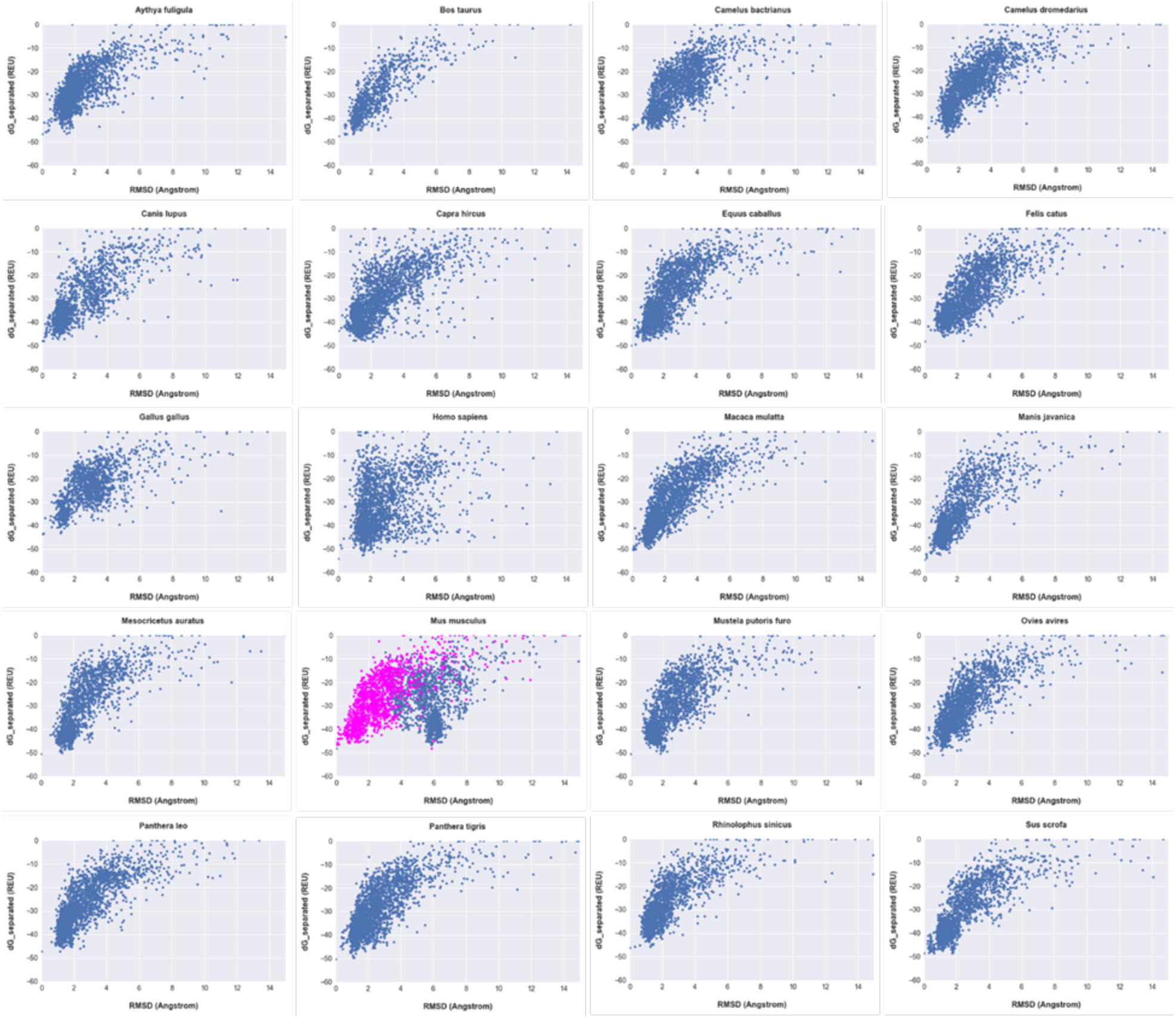
Re-docking of SARS-CoV-2-RBD to ACE2 of different species resulted in high-quality models. Cα-root mean square deviation (Cα-RMSD) were calculated against the best performing model and plotted versus predicted binding energy (dG_separated) after redocking of the SARS-CoV-2-RBD for all SARS-CoV-2-RBD-ACE2 co-complexes. This measure describes the similarity of the models compared to their predicted binding energy. Models from the lowest left corner represent the highest quality models and where chosen for further analysis. The models for *Mus musculus* were recalculated to the second-best model (magenta), as they did not converge on the best model.

**Extended Data Figure 5:**
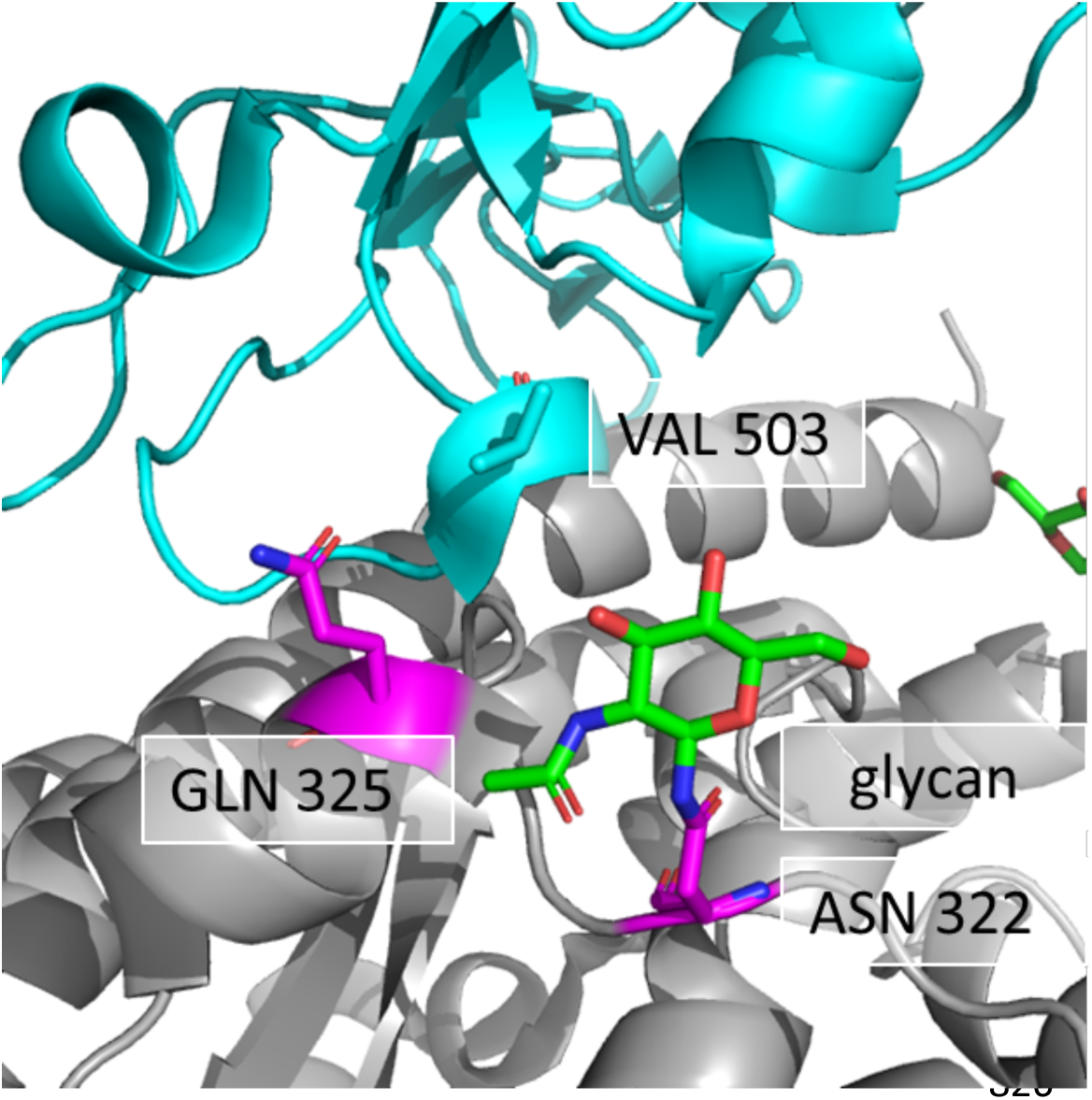
Proximity of Val503 of the SARS-CoV-2 RBD to an ACE2 glycan at Asn322. At the interface of SARS-CoV-2-RBD (cyan) and ACE2 (grey), Val503 is in close proximity to the *N-*acetylglucosamine (green) at Asn322 as seen in PDB: 6lzg.

**Extended Data Figure 6:**
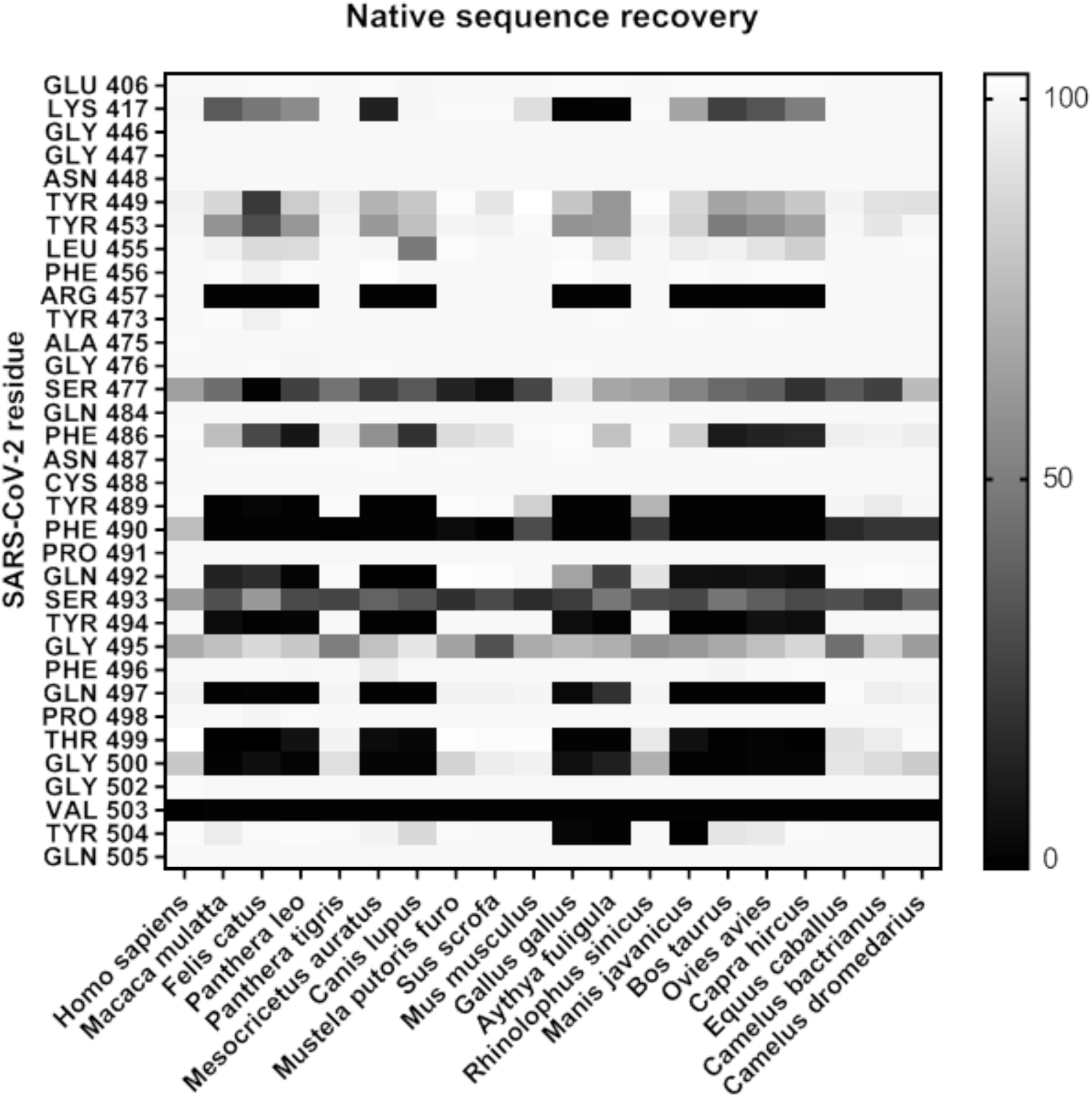
Native sequence recovery as determined by RECON multistate design allowing the mutation and optimization of SARS-CoV-2-RBD interface residues in the presence of the complex with human ACE2. This figure demonstrates all designable residues, including residues that did not change during multistate design or changed for all species in the same way.

## Supplemental Methods

### R code for calculating susceptibility scores on new species

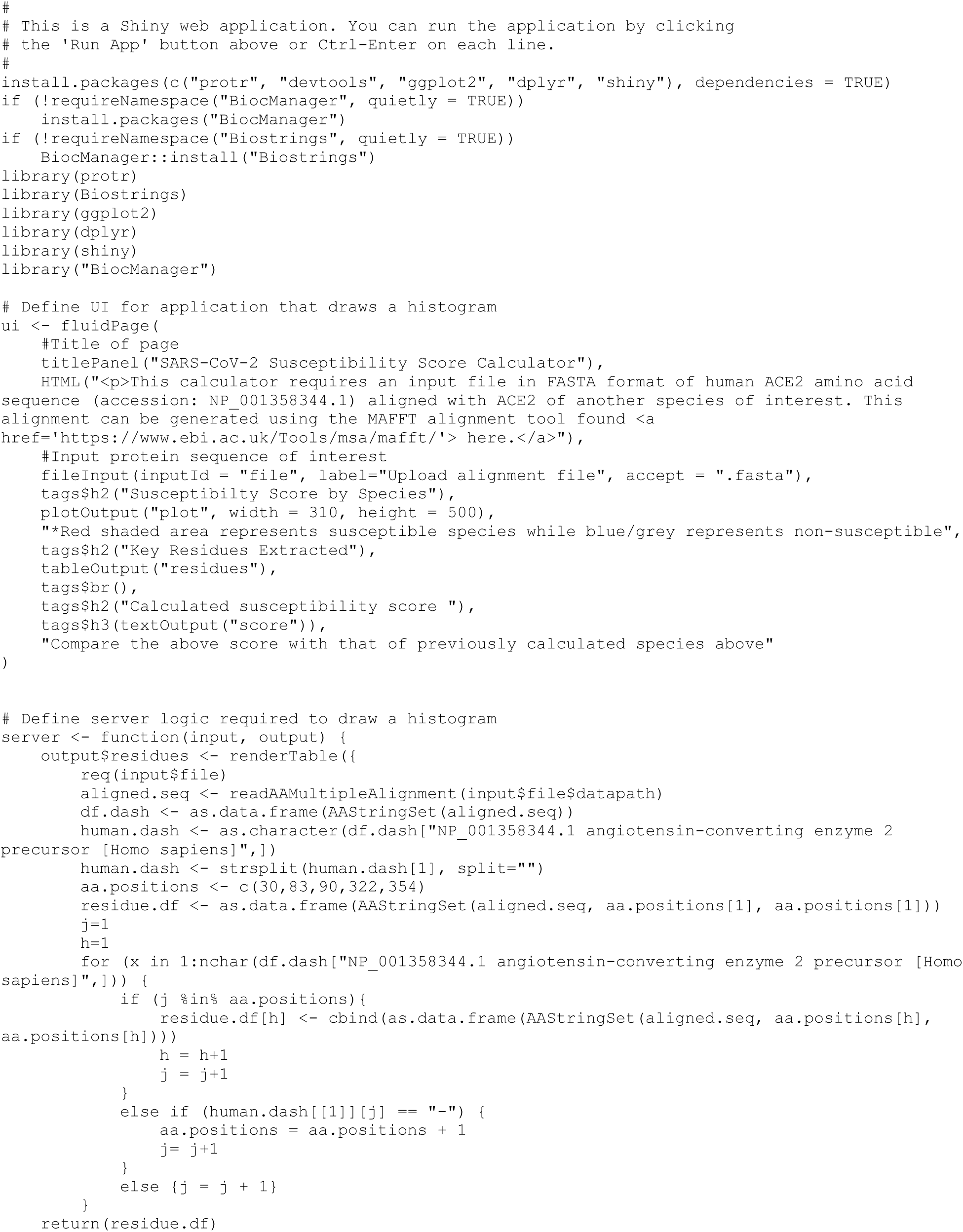

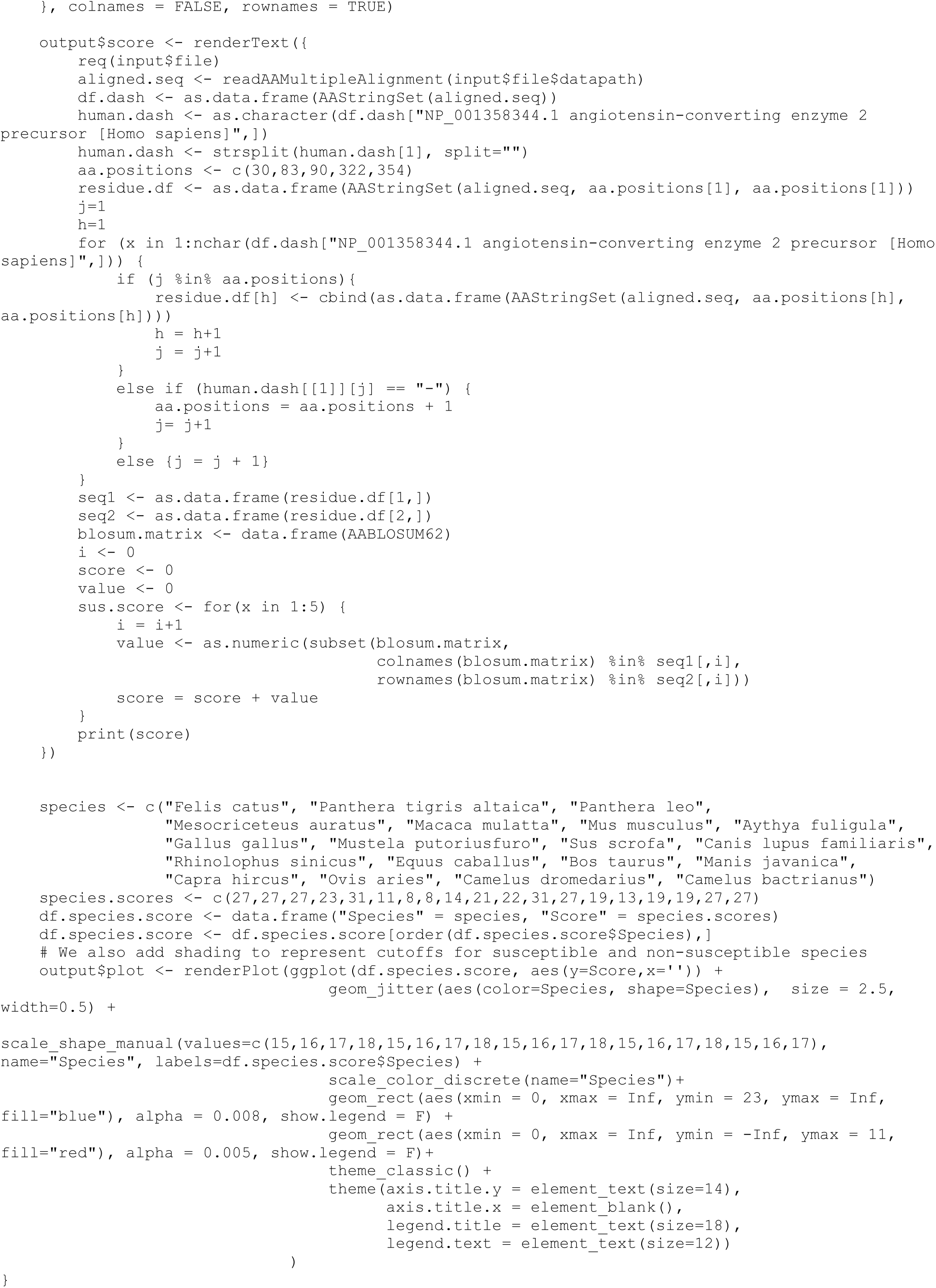

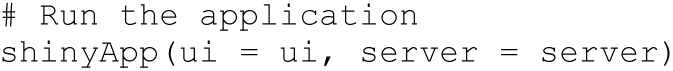

### Homology modeling of ACE2 based on the ACE2-SARS-CoV-2-RBD co-crystal structure using RosettaCM

#### Structure and input preparation

For all modeling purposes Rosetta-3.12 was used.

Preparation of input structures using RosettaRelax:

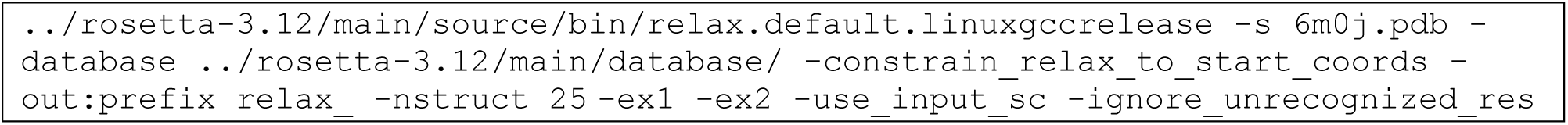

##### Command used for partial thread

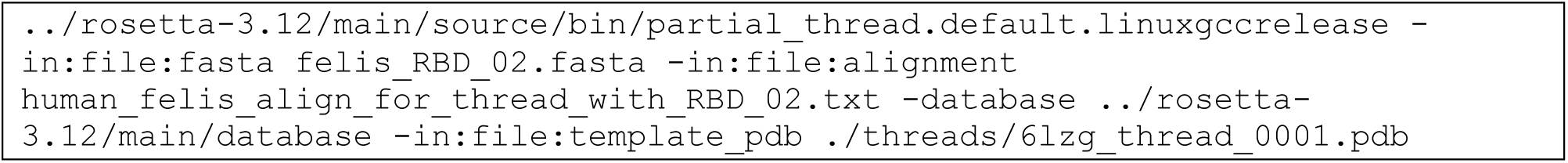

#### Construction of the initial ACE2-SARS-CoV-2-RBD complex with RosettaCM

Command used for executing RosettaCM

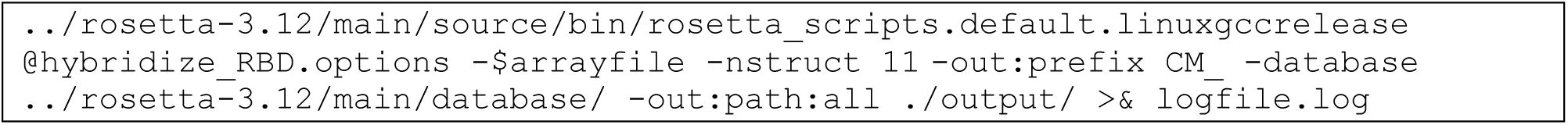

##### RosettaScripts protocol for RosettaCM

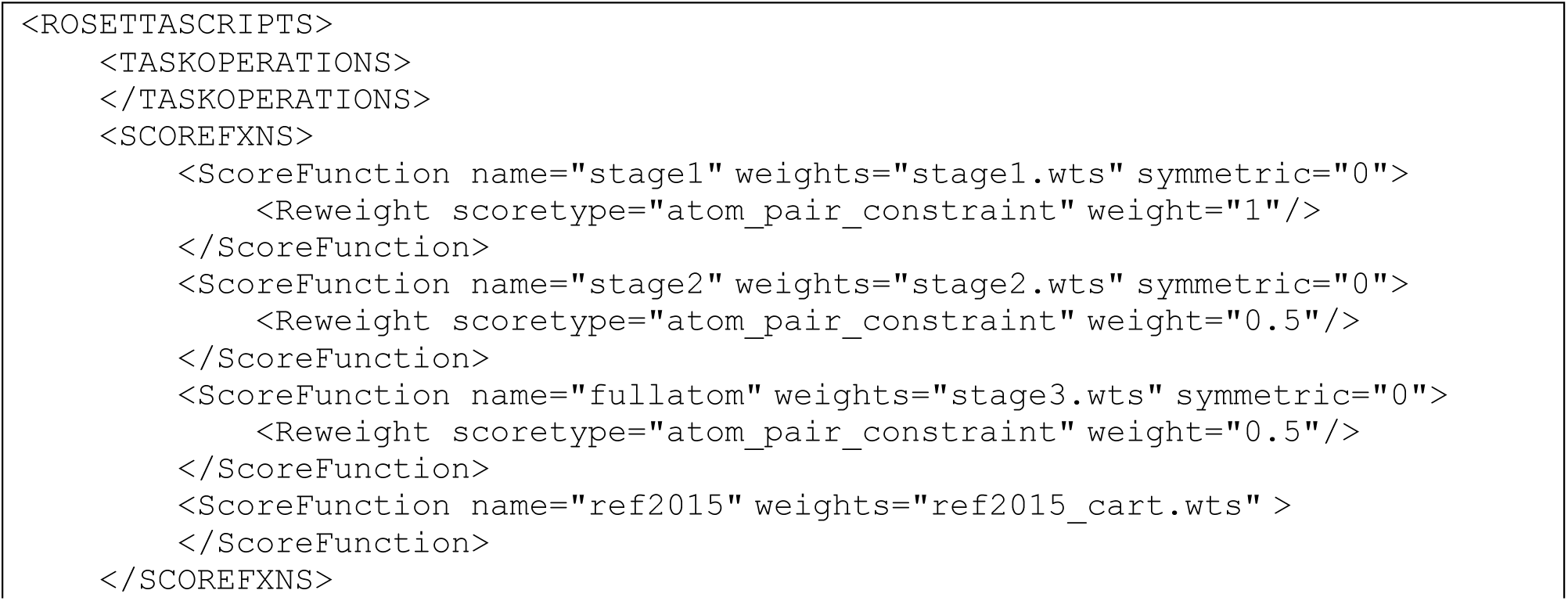

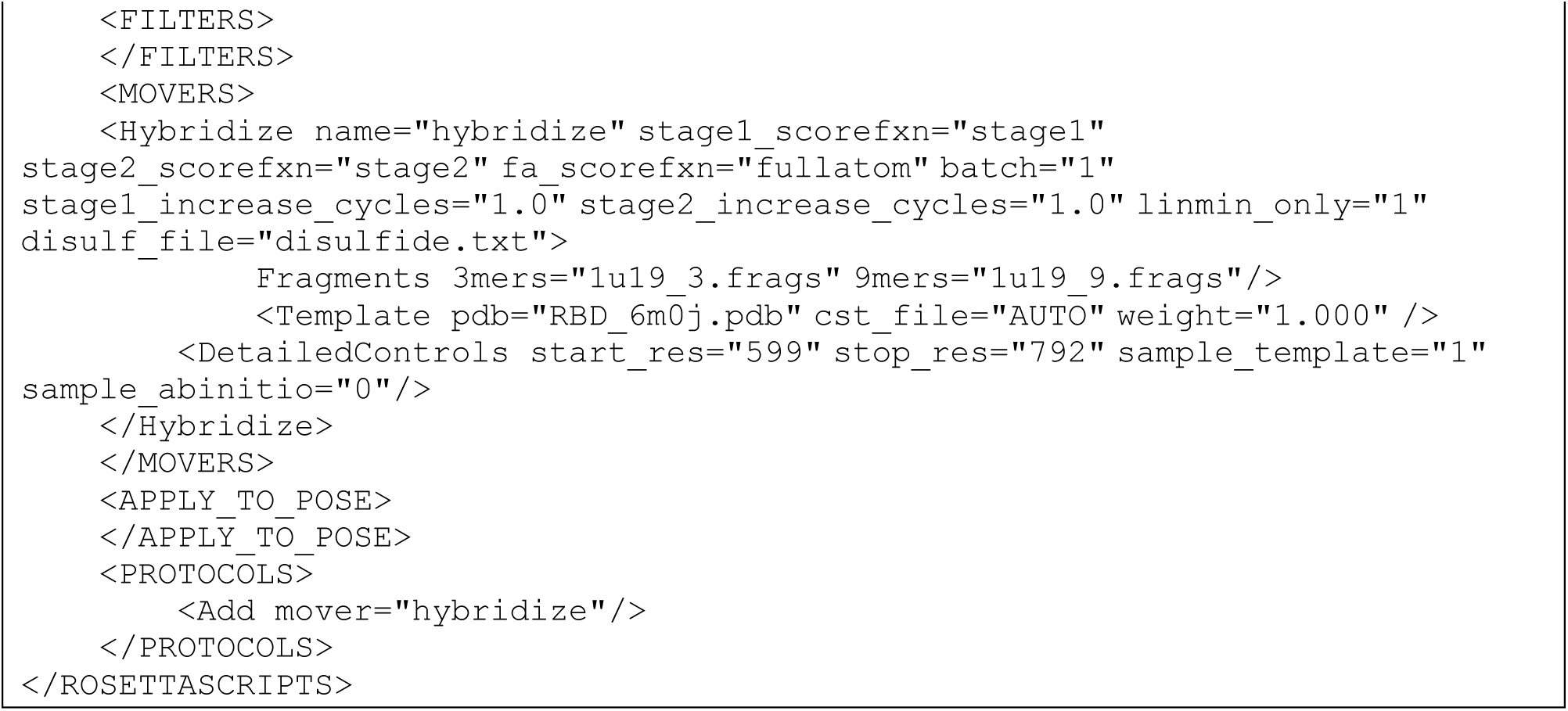

##### RosettaCM options

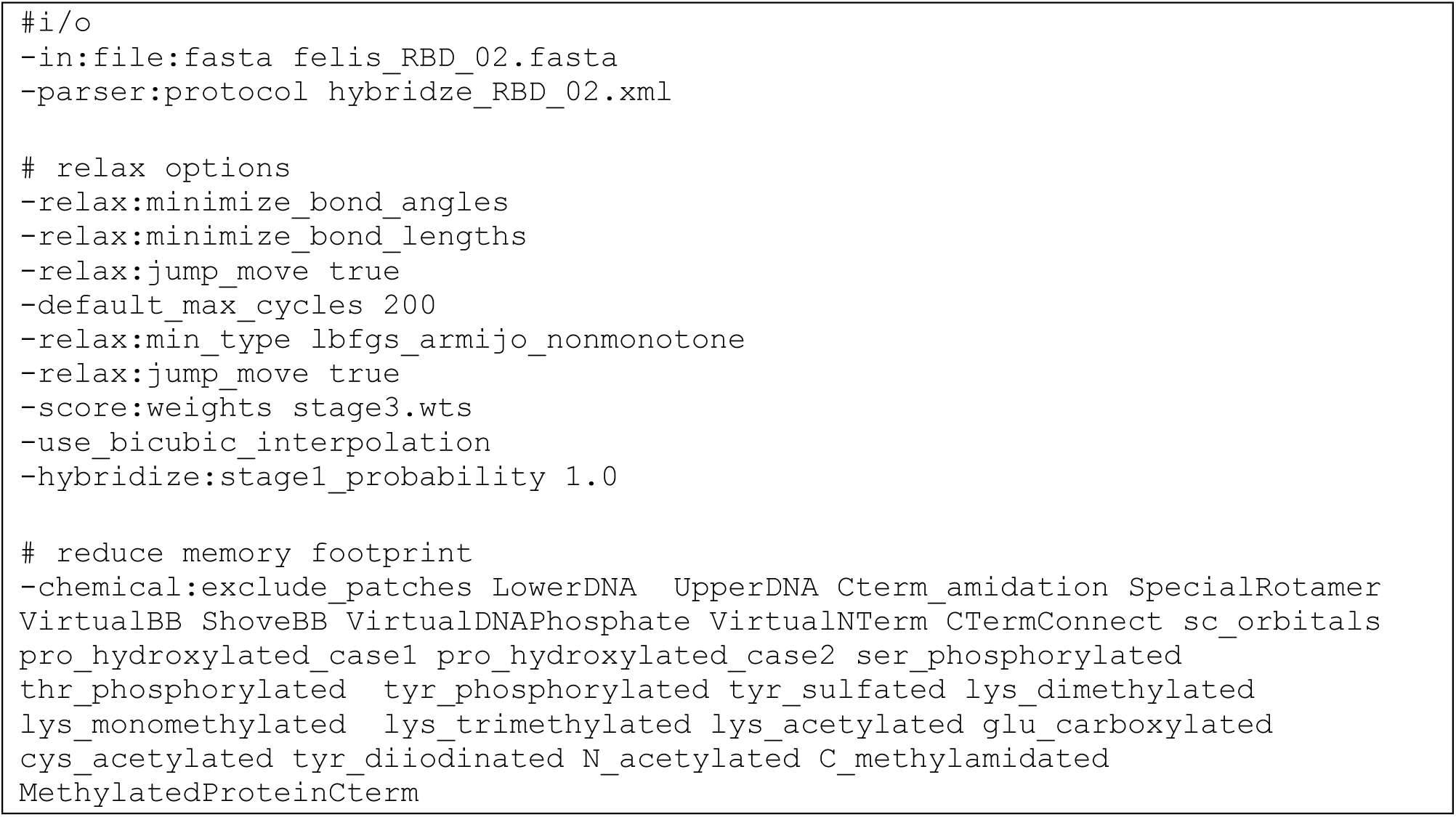

##### Stage1 weights

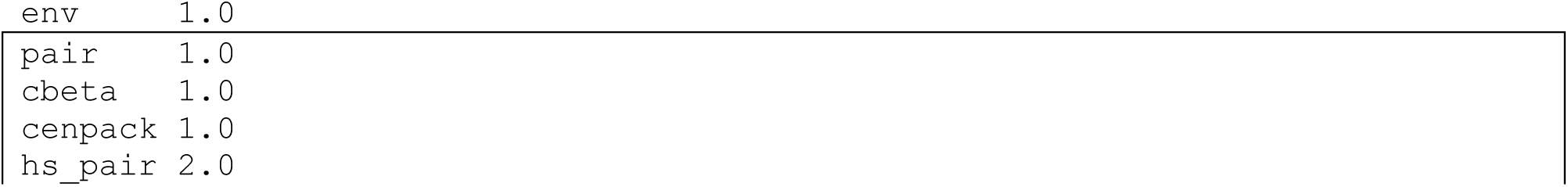

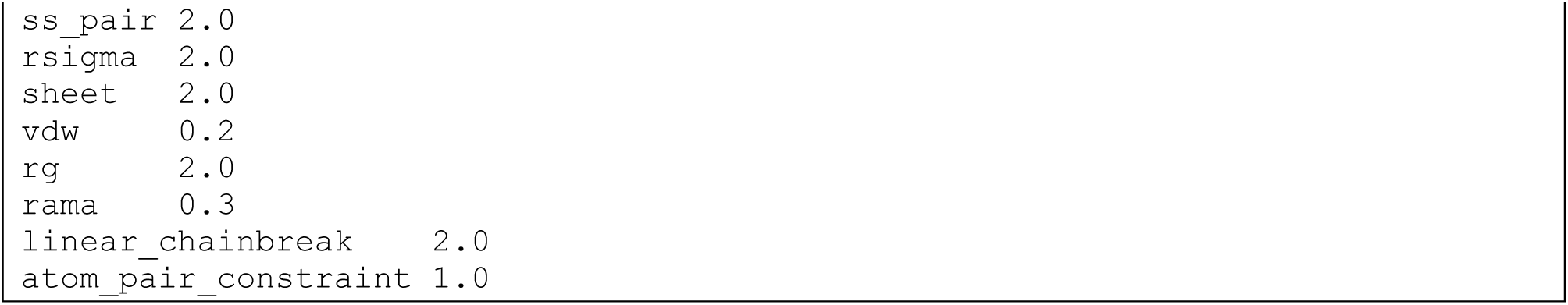

##### Stage2 weights

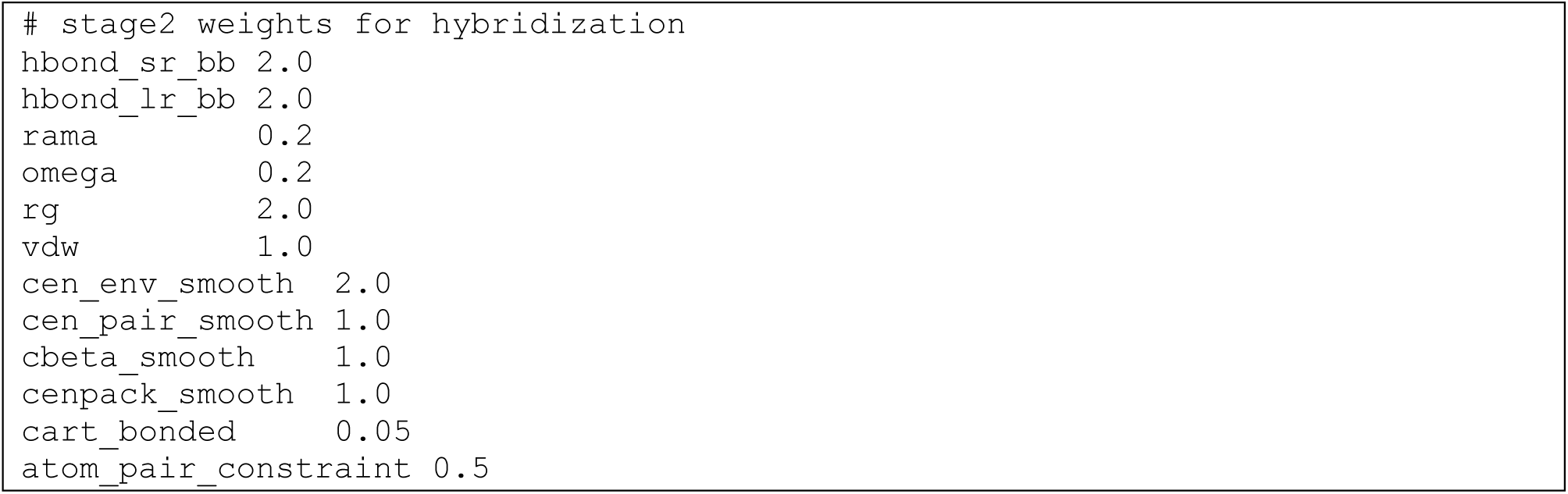

##### Stage3 weights

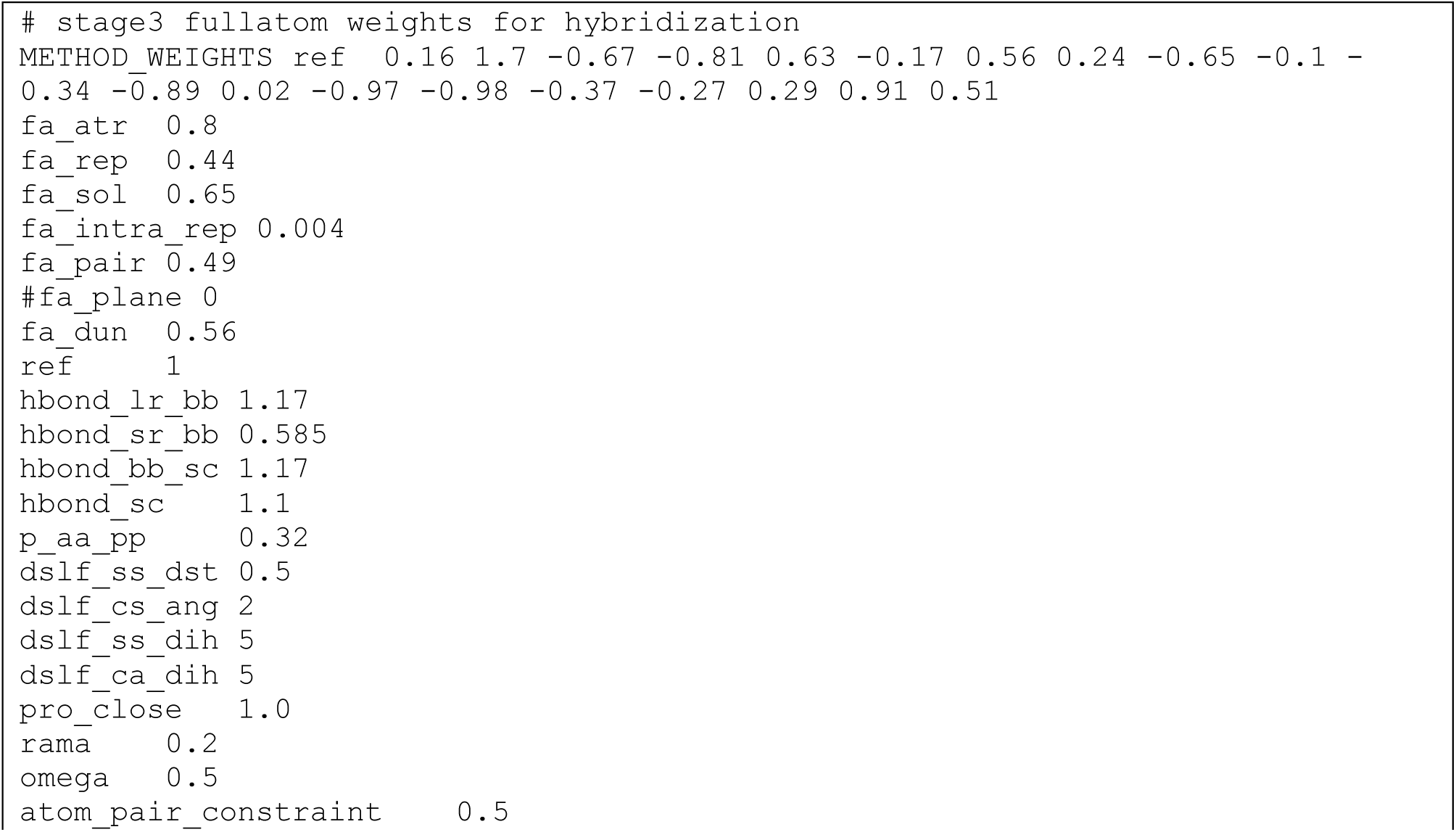

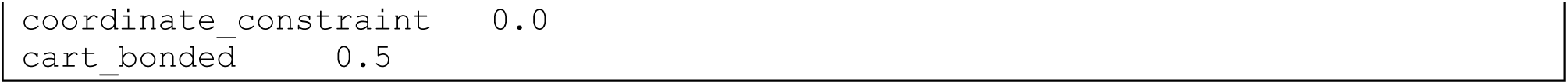

### RosettaRelax of ACE2-SARS-CoV-2 complex

#### Command for RosettaRelax

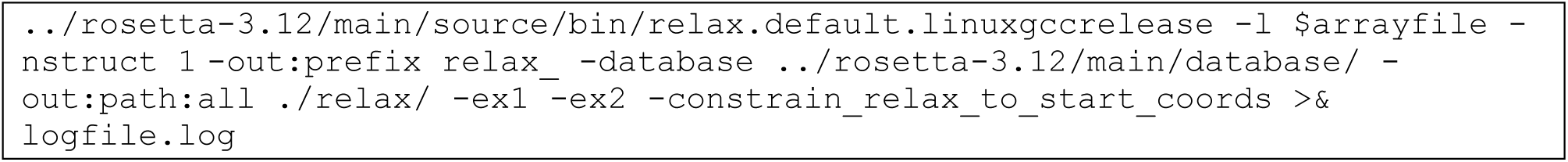

### Docking of ACE2 and SARS-CoV-2 RBD

#### Command for RosettaDock

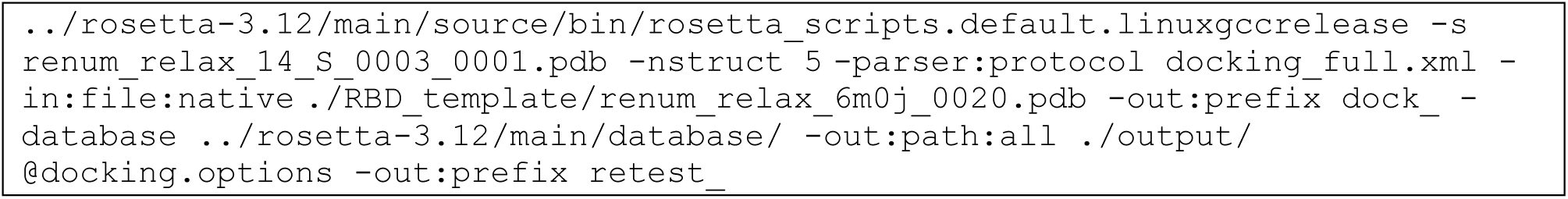

#### Options for protein-protein docking

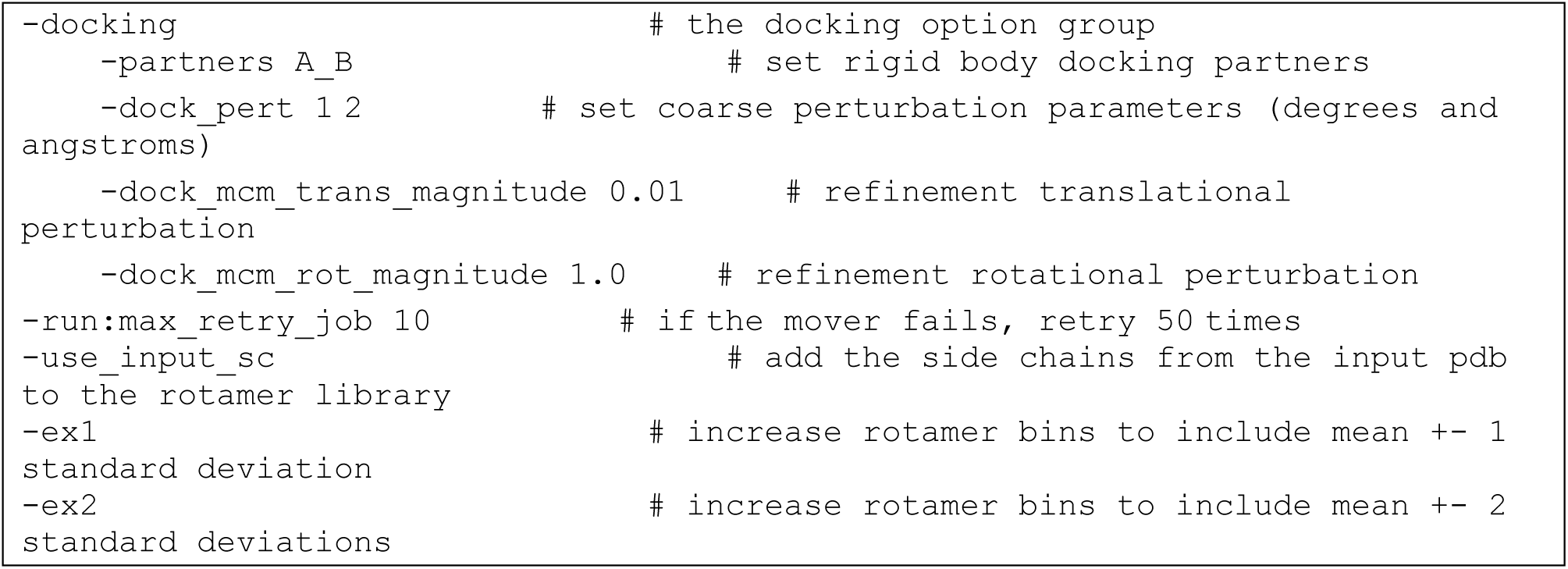

#### RosettaScripts xml-protocol for protein-protein docking

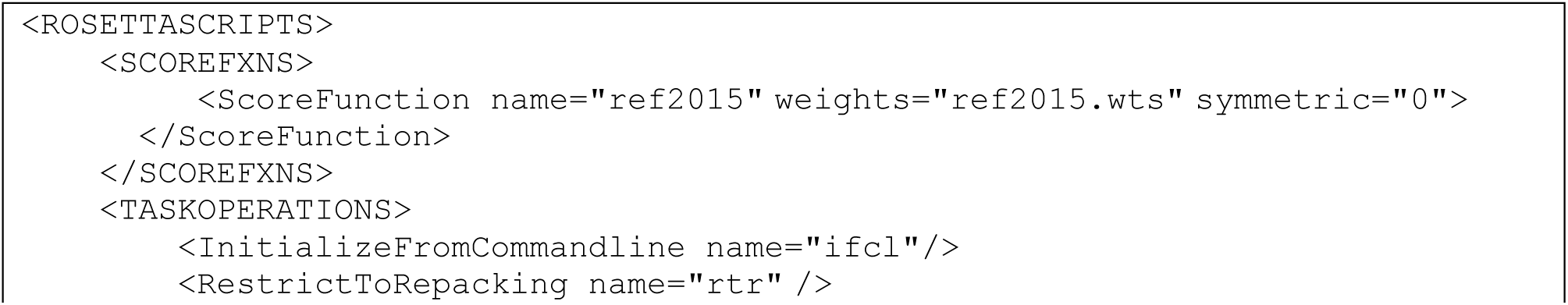

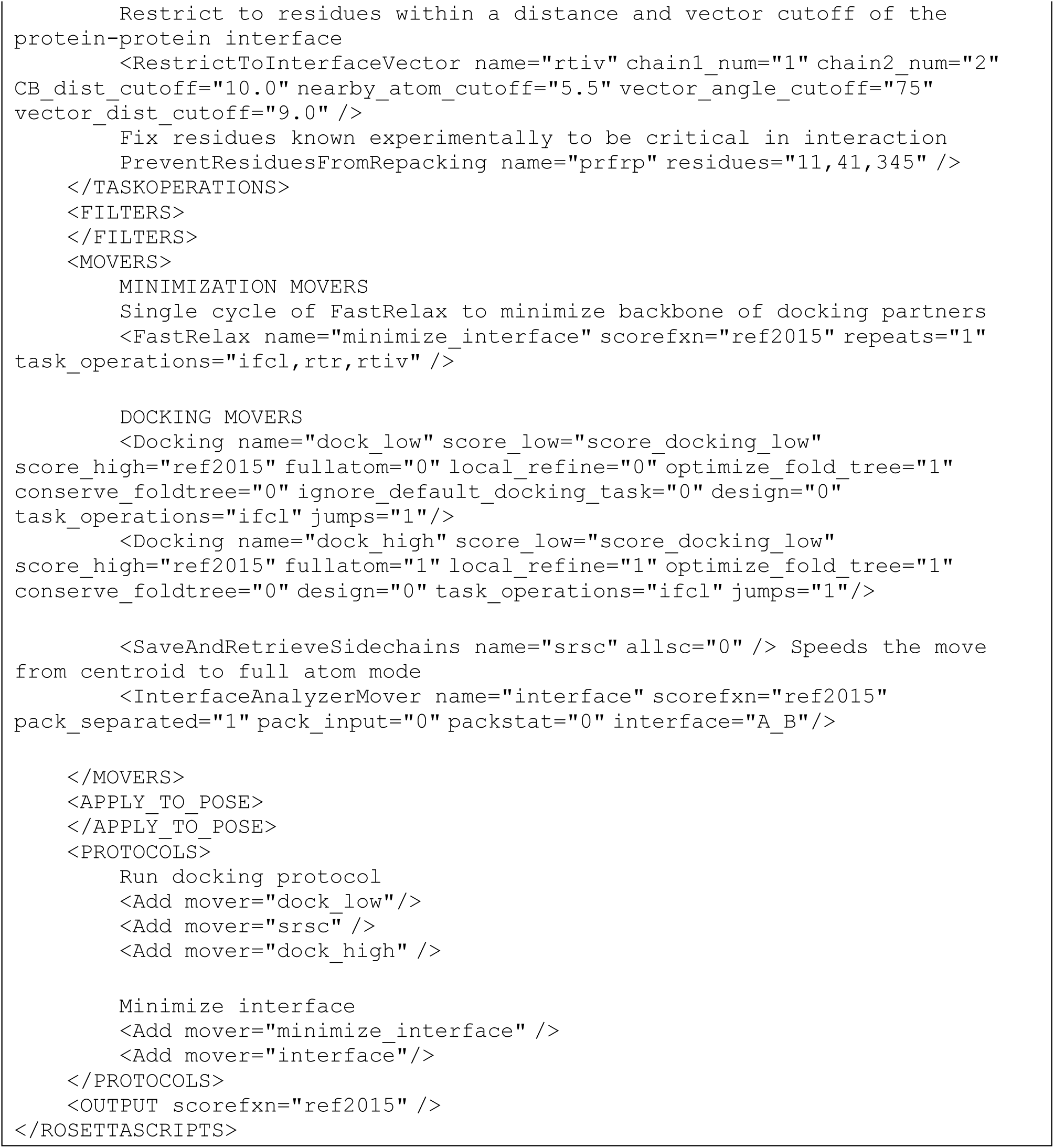

#### Options for Rosetta protein-protein docking

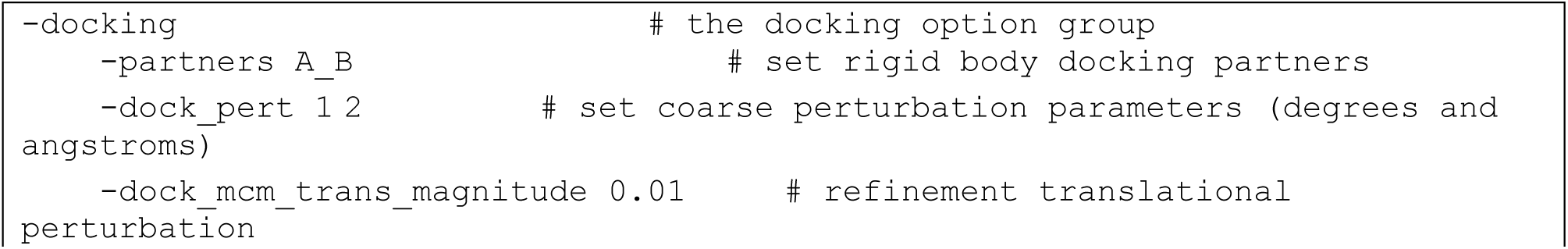

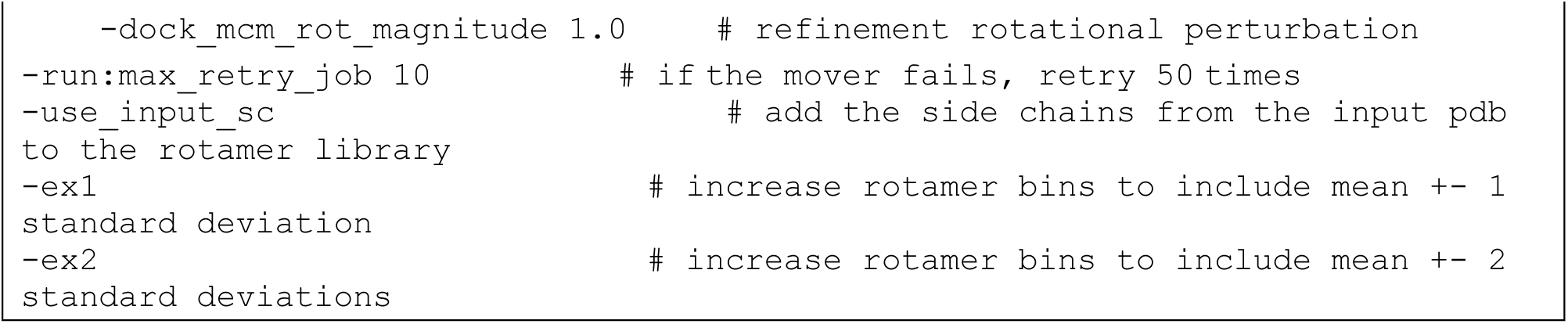

To obtain a control, the relaxed crystal structures were subjected to interface minimization.

### Calculation of native sequence recovery using the RECON multistate design protocol in Rosetta

#### Command for running RECON multistate design in Rosetta

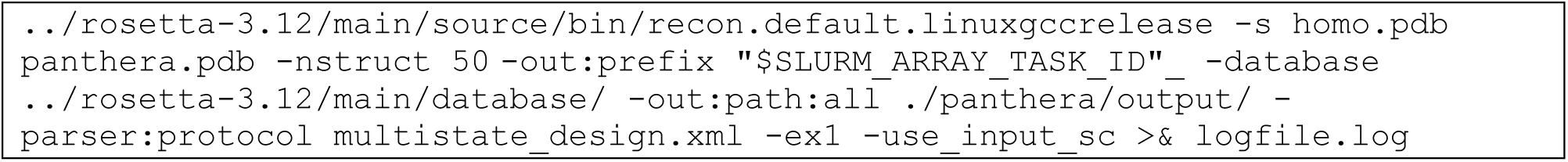

#### RosettaScripts protocol for RECON multistate design

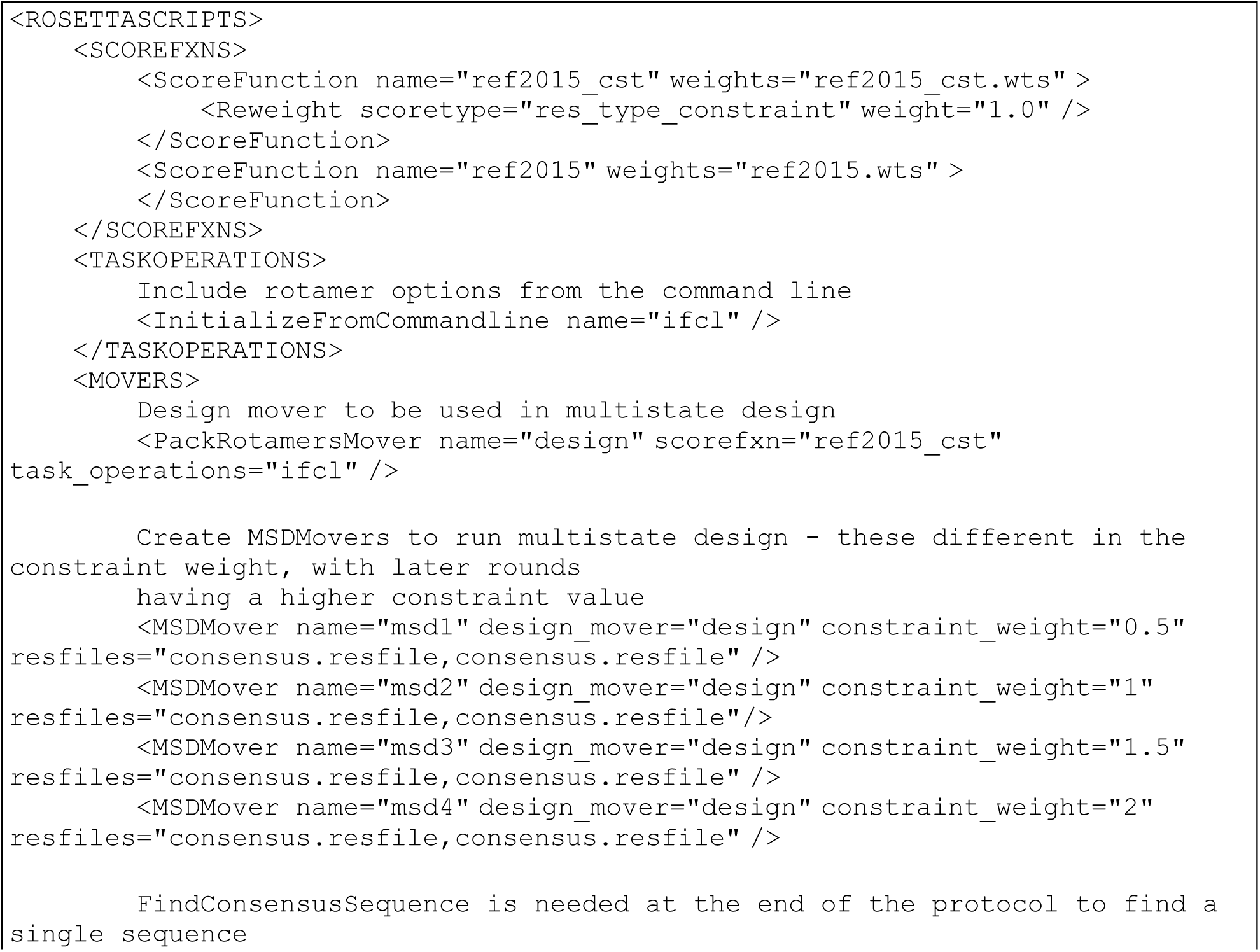

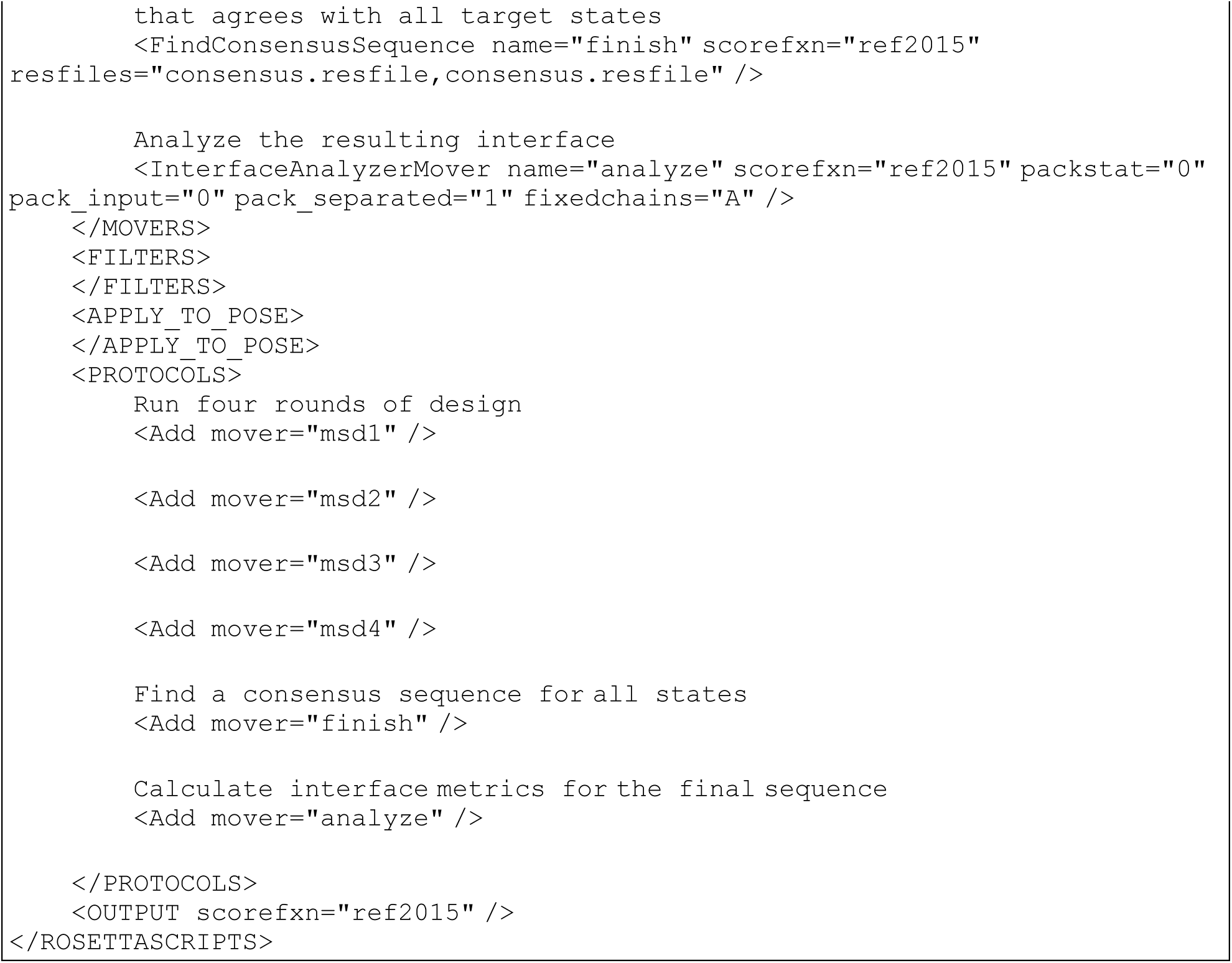

#### Calculation of native sequence recovery

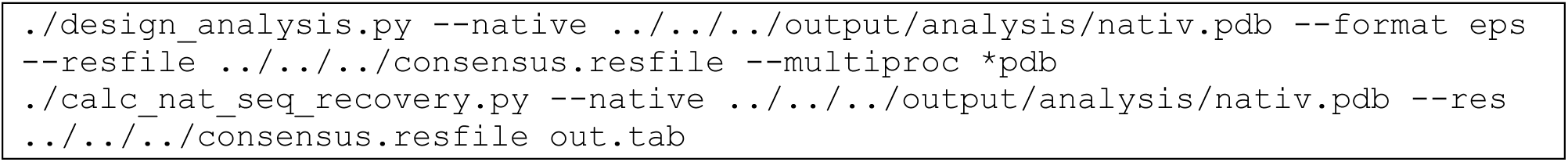

## Notes

### Competing Interest Statement

The authors have declared no competing interest.

